# Liquid-to-solid phase transition of *oskar* RNP granules is essential for their function in the *Drosophila* germline

**DOI:** 10.1101/2021.03.31.437848

**Authors:** Mainak Bose, Julia Mahamid, Anne Ephrussi

## Abstract

Asymmetric localization of *oskar* RNP granules to the oocyte posterior is crucial for abdominal patterning and germline formation of the *Drosophila* embryo. We show that *oskar* RNP granules in the oocyte are condensates with solid-like physical properties. Using purified *oskar* RNA and scaffold proteins Bruno and Hrp48, we confirm *in vitro* that *oskar* granules undergo a liquid-to-solid phase transition. Whereas the liquid phase allows RNA incorporation, the solid phase precludes incorporation of additional RNA while allowing RNA-dependent partitioning of specific proteins. Genetic modification of scaffold granule proteins, or tethering the intrinsically disordered region of human Fused in Sarcoma to *oskar* mRNA, allowed modulation of granule material properties *in vivo*. The resulting liquid-like properties impaired *oskar* localization and translation with severe consequences on embryonic development. Our study reflects how physiological phase transitions shape RNA-protein condensates to regulate localization and expression of a maternal RNA that instructs germline formation.

## Introduction

Asymmetric localization of maternal RNAs within the developing oocyte is essential for embryonic axis formation and cell fate specification in many organisms (Becalska and Gavis, 2009; Besse and Ephrussi, 2008; Buxbaum et al., 2015; Martin and Ephrussi, 2009). In *Drosophila*, *oskar* mRNA encodes the posterior determinant, Oskar protein. Posterior accumulation of Oskar is achieved by active transport of *oskar* mRNA in the form of diffraction-limited granules on a polarized microtubule network during mid-oogenesis, in a two-step transport process involving dynein and kinesin motors (Brendza et al., 2000; Clark et al., 2007; Jambor et al., 2014; Little et al., 2015; Zimyanin et al., 2008). Importantly, *oskar* mRNA is translationally repressed prior to localization, preventing ectopic production of Oskar protein (Ephrussi et al., 1991; Ephrussi and Lehmann, 1992; Kim-Ha et al., 1995). The translated Oskar protein nucleates assembly of the pole plasm, which determines abdominal patterning and germline formation in the embryo.

Inside cells, mRNAs interact with proteins to form ribonucleoprotein complexes (RNPs), in which the protein composition is dynamically remodeled during the mRNA life cycle (Moore, 2005). At high local concentrations, individual RNPs can condense into higher-order assemblies by virtue of multivalent protein-protein, RNA-protein and/or RNA-RNA interactions. These mesoscale assemblies, referred to as RNP granules, belong to the expanding class of membraneless compartments or biomolecular condensates (Tauber et al., 2020; Van Treeck and Parker, 2018), some of which form by liquid-liquid phase separation (LLPS) (Banani et al., 2017; Shin and Brangwynne, 2017). The collective behavior of RNPs in the condensed state confers emergent properties to the granules that the individual RNPs lack, and that may also evolve with time (Alberti, 2017; Alberti et al., 2019). For example, reconstitution experiments with the stress granule component Fused in Sarcoma (FUS) have shown that FUS droplets assemble by LLPS into spherical condensates that mature with time into a solid non-dynamic state, a phenomenon described as ageing (Patel et al., 2015). Condensate ageing is physiologically pertinent as evident from the *C. elegans* peri-centriolar matrix (PCM), which exhibits distinct liquid-like and solid-like physical states at different stages of the embryonic cell cycle (Mittasch et al., 2020; Woodruff et al., 2017; Woodruff et al., 2018). Thus, cells can harness different condensate properties to achieve specific functions.

We report that *oskar* granules are RNA-protein condensates that behave like solids *in vivo* and *in vitro*, unlike the vast majority of liquid-like RNP granules described to date (Banani et al., 2017; Brangwynne et al., 2009; Brangwynne et al., 2011; Fujioka et al., 2020; Shin and Brangwynne, 2017; Wippich et al., 2013). *In vitro* reconstitution of *oskar* 3’UTR with scaffold granule proteins lead to formation of amorphous, spherical and dynamic condensates that rapidly mature into a solid state. We demonstrate that this liquid-to-solid transition is physiologically essential, as perturbing the solid state *in vivo* impaired RNA localization and translation, linking regulation of granule material properties to RNA post-transcriptional control.

## Results

### *oskar* RNP granules in the oocyte are spherical solid-like compartments

Proteins involved in transport and/or translational control associate with *oskar* mRNA. The *bona fide* RNP components Bruno and PTB have been shown to oligomerize on the *oskar* 3’UTR, forming higher-order complexes (Besse et al., 2009; Chekulaeva et al., 2006; Kim et al., 2015). *oskar* has also been reported to dimerize by virtue of a stem-loop structure in the 3’UTR (Jambor et al., 2011). These findings suggest that multivalent interactions between *oskar* and associated proteins promote formation of higher-order transport granules and prompted us to investigate the potential role of biomolecular condensation in *oskar* granule assembly and function. A liquid condensate assumes a spherical shape due to surface tension (Widom, 1988). This shape criterion determined by observations made with light microscopy has been central in assessing whether granules assemble via LLPS (Alberti et al., 2019; Hyman et al., 2014). However, *oskar* granules in the oocyte are diffraction-limited point sources, which precluded characterization of their shape by conventional confocal microscopy. We therefore resorted to 3D STED (stimulated emission depletion) super resolution imaging with near isotropic resolution. We unequivocally show that *oskar* granules are spherical, with an aspect ratio ∼1 and a Gaussian distribution of sizes (Figure 1A and Movie S1). This is consistent with the notion that assembly *in vivo* is driven by LLPS.

**Figure 1.**
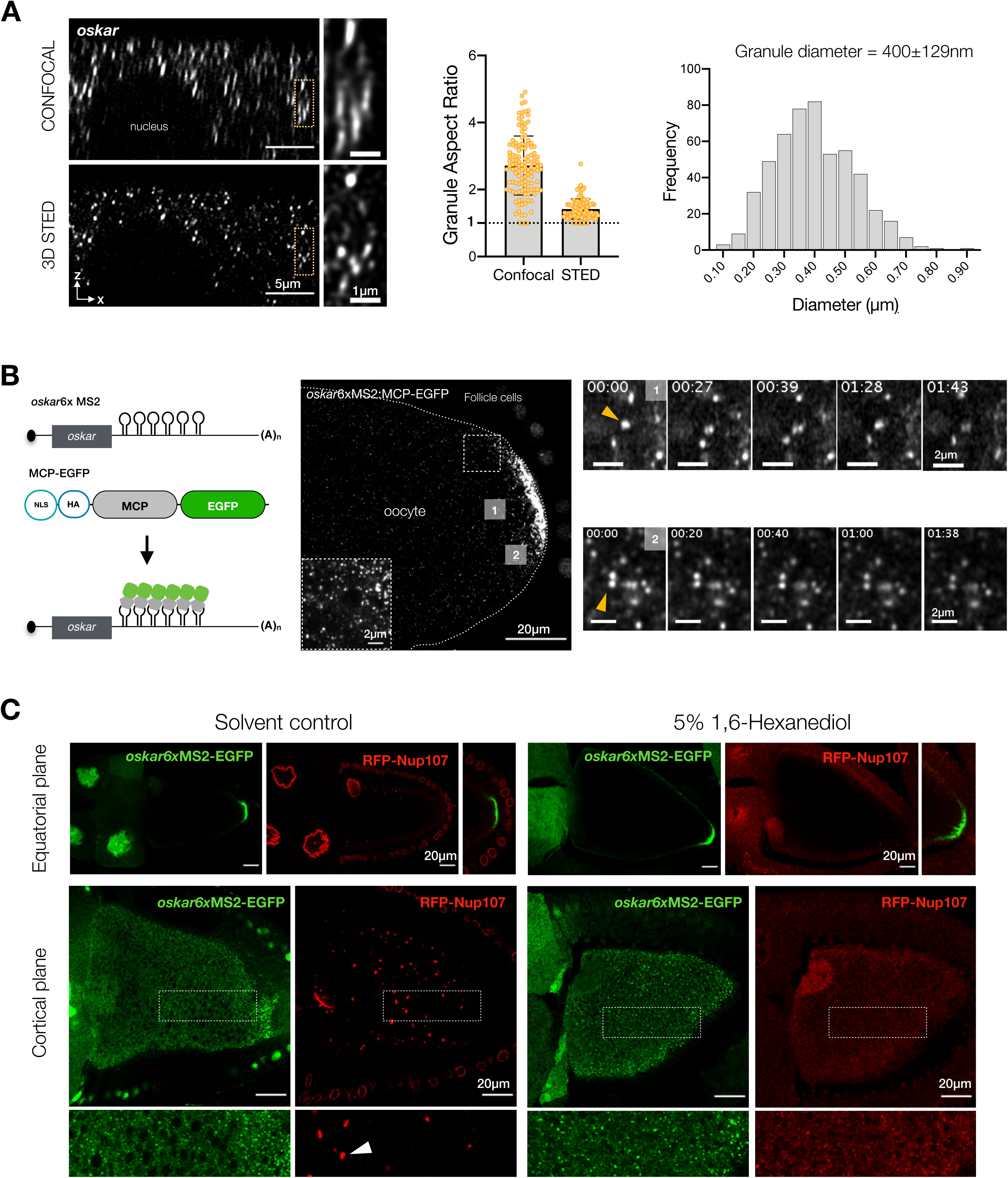
*oskar* RNP granules are spherical, solid-like assemblies. (A) Comparison of confocal and 3D STED channels in XZ plane, with *oskar* (grayscale) detected by single-molecule fluorescence *in situ* hybridization (smFISH) in wild type (*w^1118^*) egg chambers. Comparison of granule aspect ratio, and diameter distribution as measured from STED volume imaging. 3D volumes were acquired with a voxel size of 40nm x 40nm x 60nm. Error bars represent SD. (B) Schematic representation of *oskar* mRNA *in vivo* labelling strategy using the MCP-MS2 system. A representative stage 10 egg chamber showing *oskar* mRNA labelled with MCP-EGFP. Boxed area is enlarged at bottom left. Live imaging of MCP-EGFP tethered *oskar* granules was performed on a cortical region approximately 15-20 µm from the posterior pole (1) and on granules at the posterior pole (2). The time-point (min:sec) is indicated at the top of each frame. (C) Representative images of egg chambers expressing RFP-Nup107 and *oskar*6xMS2-MCP-EGFP after treatment with solvent control or 5% 1,6-hexanediol for 15 min. Neither *oskar* granules at posterior pole (equatorial plane imaging) nor individual granules (cortical plane imaging) are dissolved by hexanediol, whereas nuclear envelopes and annulate lamellae (white arrowhead) are dissolved. Boxed areas are enlarged at the bottom.

Upon contact, two liquid-like condensates typically fuse and rearrange into a single spherical structure (Brangwynne et al., 2009; Hyman et al., 2014). To investigate the dynamic behavior of *oskar* RNP granules, we tagged *oskar* mRNA with enhanced green fluorescent protein (EGFP) using the MCP-MS2 tethering system (Figure 1B). Imaging in live egg chambers near the cortical surface visualized occasional directed transport on microtubule tracks, in addition to diffusive movements due to Brownian motion and cytoplasmic flows. Interestingly, two granules that touched each other did not fuse and relax into one within the timescale of imaging (4 minutes, Figure 1B (1); Movie S2). At the posterior pole, granules that are presumably anchored also did not fuse despite their high local concentration, indicating that *oskar* granules are not liquid-like (Figure 1B (2); Movie S3).

A liquid phase is susceptible to dissolution upon dilution (Putnam et al., 2019). Extrusion of the ooplasm into a physiological buffer does not result in dissolution of *oskar* granules, further confirming their non-liquid properties (Gaspar and Ephrussi, 2017; Gaspar et al., 2017). Furthermore, treatment of egg chambers with 1,6-hexanediol, a chemical that may perturb weak multivalent interactions in LLPS and is employed as a probe to distinguish liquid from solid condensates (Kroschwald et al., 2017), dissolved phase-separated precursors of nuclear envelopes labelled with red fluorescent protein (RFP)-Nup-107 (Hampoelz et al., 2019), but had no effect on *oskar* granules (Figure 1C). Taken together, these observations indicate that *oskar* RNP granules *in vivo* are phase separated solid-like assemblies.

### *Bona fide oskar* granule proteins are RNA binding proteins with structural disorder

Genetic studies identified several RNA binding proteins (RBPs) that associate with *oskar* mRNA and engage in diverse processes, including RNP transport and translation repression. However, due to the syncytial architecture of the germline, biochemical experiments lack spatial information limiting knowledge about when and where within the egg chamber, and at what stoichiometries, these proteins associate. The proteins Bruno, PTB (Polypyrimidine Tract Binding protein) and Hrp48 bind specific sequences along the RNA and regulate its translation. (Besse et al., 2009; Huynh et al., 2004; Kim-Ha et al., 1995; Snee et al., 2008; Yano et al., 2004). Additionally, Bruno and PTB have been implicated in formation of higher-order structures with the *oskar* 3’UTR (Besse et al., 2009; Chekulaeva et al., 2006). We therefore hypothesized that these *bona fide* granule components may form the scaffold of the granule and contribute to condensation.

We used imaging to determine where the candidate proteins associate with *oskar* (Figure 2A and S1A). Bruno associated with *oskar* in the nurse cell cytoplasm and colocalized with *oskar* mRNA on track-like structures. The association is maintained in the ooplasm. Hrp48 was diffuse within the nurse cell cytoplasm, with occasional enrichment with *oskar* on track-like structures. Granular appearance and *oskar* association of Hrp48 became prominent in the ooplasm. PTB, on the other hand was largely nuclear in the nurse cells with no obvious colocalization with *oskar,* but associated with *oskar* in the oocyte, indicating sequential recruitment of proteins to *oskar* granules.

**Figure 2.**
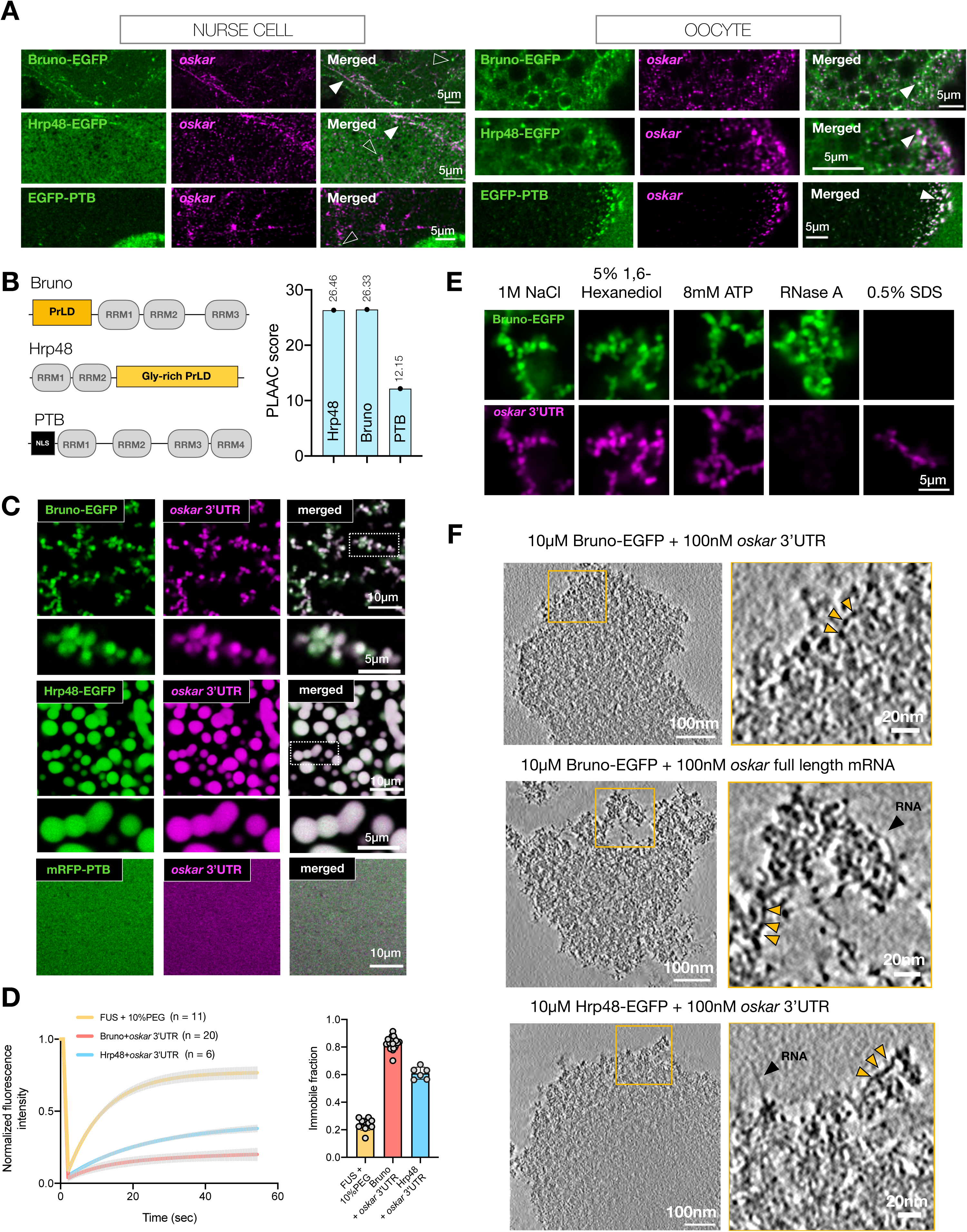
*In vitro* reconstituted minimal *oskar* RNP condensates recapitulate properties of *in vivo* RNP granules. (A) *oskar* mRNA association with the three RBPs in the nurse cell cytoplasm and oocyte posterior (cortical plane imaged). A maximum intensity projection of a Z-stack of 1 µm with *oskar* mRNA smFISH (magenta) and the respective proteins (green). White arrowheads indicate colocalization. Empty arrowheads represent foci where no colocalization is detected. Refer to Figure S1A for full images. (B) Domain architecture of 3 *oskar* granule component proteins and their PLAAC score. (C) Condensates formed with 100 nM *oskar* 3’UTR-atto633 (magenta) and 10 µM of the indicated RBPs (green) imaged with confocal microscopy. Framed areas are enlarged at the bottom. (D) Quantification of fluorescent recovery after photobleaching (FRAP) kinetics and immobile fraction of condensates assembled with 100 nM *oskar* 3’UTR (unlabeled) and 10 µM of Bruno-EGFP (green) or Hrp48-EGFP (green). hFUS-EGFP condensates as control were assembled with 8 µM hFUS-EGFP (without RNA) and 10% PEG-4000 (Patel et al., 2015). Error bars represent SD. *n* denotes number of movies per condition. (E) Bruno-EGFP (green)-*oskar* 3’UTR (magenta) condensates subjected to the indicated treatments after 30 min of ageing. Images acquired under identical microscope settings. (F) Tomographic slices, 4 nm thick, of condensates formed under the indicated conditions and plunge-frozen after 30 min of ageing. Insets are marked with yellow boxes. Condensates are largely amorphous with frequent ‘beads on a string’ structures (yellow arrowheads). Putative naked RNA molecules are marked with black arrowheads. For full tomogram volume refer to movies S8-S10. See also figures S1, S2 and S3.

RBPs with prion-like domains (PrLDs) play key roles in RNP granule formation by promoting multivalent interactions involving RNA, folded protein domains, PrLDs of their own and/or of other proteins (Protter et al., 2018). We asked if *oskar*-binding RBPs possess unstructured domains. Using the prion prediction algorithm PLAAC (Lancaster et al., 2014), we identified that Bruno has an N-terminal PrLD with an over representation of Ser and Asp. Scoring similar to Bruno, Hrp48 possesses a 200 residue-long PrLD enriched in Ser and Gly. PTB, which consists of 4 RRMs (RNA Recognition Motif), lacks domains of substantial disorder (Figure 2B and S1B). The *oskar* granule proteins Bruno and Hrp48 may therefore be sufficiently disordered to drive LLPS, whereas PTB is not likely to directly contribute to condensation.

### *In vitro* reconstituted minimal *oskar* RNP condensates undergo liquid to solid phase transition

We purified EGFP-tagged full length Bruno and Hrp48 from insect cells using a solubility tag (6xHis-SumoStar) (Figure S2A). Electrophoretic mobility shift assay (EMSA) confirmed that all three proteins directly bind *oskar* 3’UTR *in vitro* (Figure S2B). Cleavage of the solubility tag coupled with a buffer exchange to physiological salt concentration (150 mM NaCl) triggered self-assembly of both proteins into spherical condensates (Figure S2C-D). Condensation of Bruno and Hrp48 was also observed in the presence of *in vitro* transcribed *oskar* 3’UTR (Figure 2C). Bruno-*oskar* 3’UTR condensates were ∼2 µm in diameter and tended to stick to each other, while Hrp48 formed larger droplets with *oskar* 3’UTR. Notably, under the same conditions *oskar* 3’UTR alone did not self-assemble (Figure S2E). Monomeric red fluorescent protein (mRFP)-PTB was soluble (Figure 2C) and only formed condensates in the presence of a crowding agent, in the presence or absence of *oskar* 3’UTR (Figure S2F).

We did not observe fusion events when *oskar* 3’UTR condensates with Bruno or Hrp48 settled on the surface of the glass (Figure S2G, Movies S4 and S5). In contrast, condensates formed by hFUS-EGFP in the same experimental setup fused as reported previously (Movie S6) (Patel et al., 2015). Unlike FUS, recovery from photobleaching was negligible when measured within Bruno-*oskar* 3’UTR condensates. Hrp48 condensates showed intermediate recovery kinetics (Figure 2D, S2H, Movie S7). The spherical shape, but lack of fusion and FRAP recovery, indicates a rapid liquid-to-solid phase transition *in vitro*, with Bruno showing a larger immobile fraction compared to Hrp48. Owing to the diffraction-limited size and the high local abundance at the posterior pole, FRAP on *oskar* granules *in vivo* could not be performed (Kistler et al., 2018). Bruno-*oskar* 3’UTR condensates were stable under conditions that promote dissolution of liquid droplets (Figure 2E) (Putnam et al., 2019). 0.5% SDS dissolved Bruno from the condensates, confirming that Bruno does not transition into amyloids but forms a stable solid-like phase. The signal from the residual RNA (Figure 2E) indicates the formation of RNA-RNA interactions within the condensate induced upon Bruno-driven condensation, as *oskar* 3’UTR alone did not self-assemble under identical conditions (Figure S2E). Cryo-electron tomography (cryo-ET) confirmed that *in vitro* condensates which are below the diffraction-limit of a conventional light microscope are spherical in shape (Figure S3A-B). Cryo-ET of the protein condensates either in the presence of 3’UTR or the full length *oskar* mRNA showed that the condensates appear amorphous and confirmed that a liquid-to-solid phase transition was not associated with structural rearrangement of the component molecules (Figure 2F, S3, Movies S8-S10) (Jawerth et al., 2020).

The observed material properties of the minimal *in vitro* condensates are reminiscent of the solid-like nature of *oskar* granules in the oocyte, and suggest that a liquid-to-solid phase transition follows *oskar* RNP assembly *in vivo*. We could not detect a liquid-like state of the granules *in vivo*. It is possible that rapid hardening to a solid state *in vivo* arrests the granules as sub-micron particles precluding fusion into micron-scale assemblies.

### The liquid phase is essential for incorporation of *oskar* mRNA *in vitro*

Condensation might allow packaging of several *oskar* RNA molecules within a granule for efficient localization to the oocyte posterior. Therefore, we hypothesized that a transient liquid state is required for *oskar* RNA incorporation. To test this, we assembled Bruno and *oskar* 3’UTR (100:1 molar ratio) condensates *in vitro*, then added fluorescently labelled *oskar* 3’UTR. When labelled RNA was added at 0 min, its fluorescent signal overlapped with the spherical Bruno condensates. Fluorescent RNA added after 30 min instead formed a shell on the condensate surface (Figure 3A). Lowering the ratio of protein to RNA concentrations to 20:1, to recapitulate stoichiometries closer to those we measured in the oocytes, resulted in similar exclusion of labelled RNA at 30 min (Figure S4A-C). Aged Bruno condensates formed in absence of RNA also excluded RNA added at 30 min, confirming that the exclusion is a consequence of the physical properties of the protein in an aged condensed state and not due to charge-based repulsion (Figure S4D). Cryo-ET visualized this RNA exclusion at 30 min as abundant naked RNA molecules at the periphery of the amorphous condensed phase (Figure 3B, Movies S11-12). Similar co-condensation and exclusion of RNA was observed for Hrp48 (Figure 3A).

**Figure 3.**
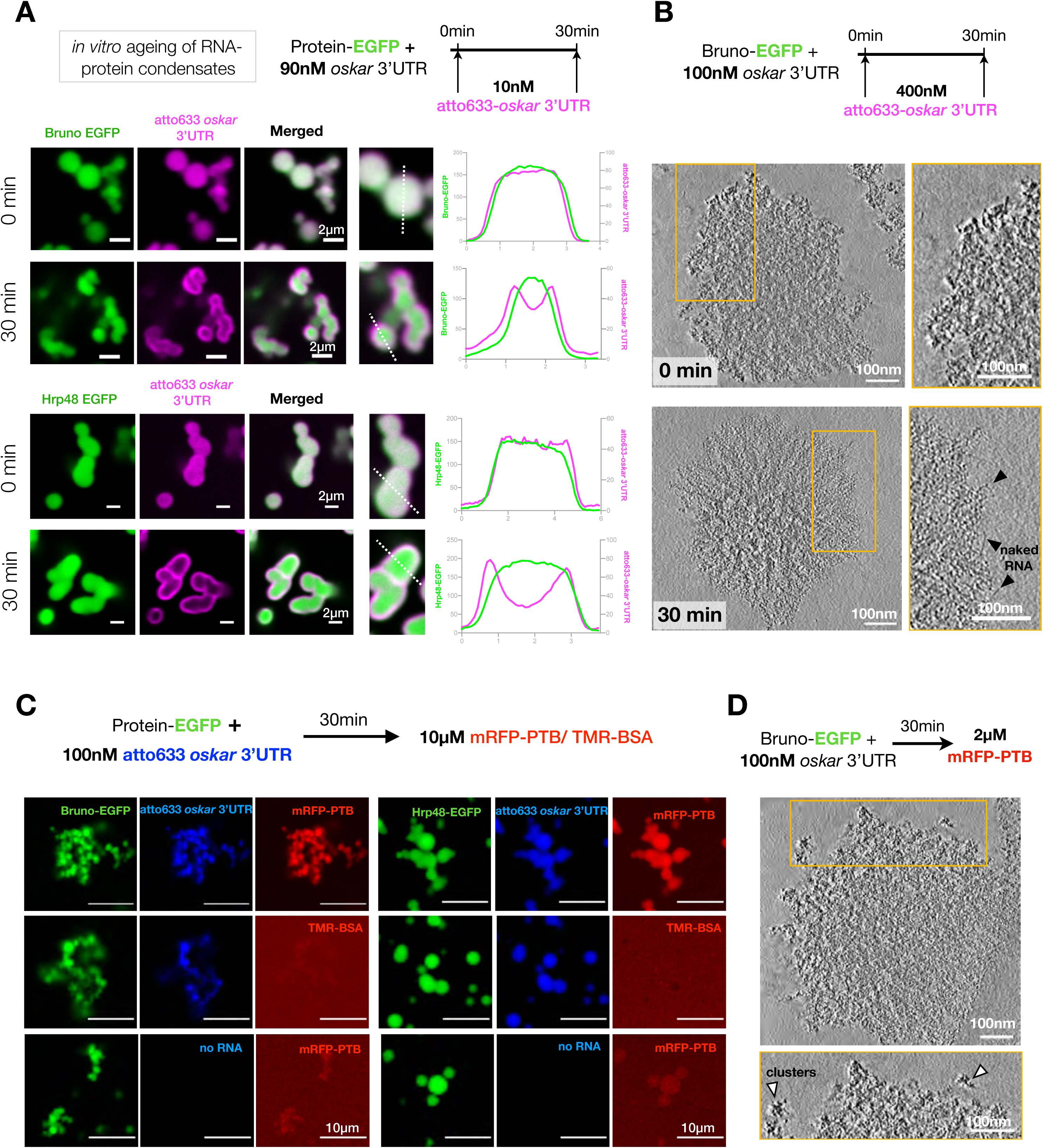
*in vitro* reconstituted *oskar* RNP condensates are selectively permeable. (A) Scheme of the *in vitro* condensate ageing assay. Single confocal slices and representative line profiles (along dotted line) shown. (B) 400 nM atto633-*oskar* 3’UTR was added at 0 and 30 min to Bruno condensates assembled with 100 nM *oskar* 3’UTR (unlabeled) as depicted in the schematic. Light microscopy images in Figure S4C. Tomographic slices (4 nm thick) of condensates at 0 and 30 min after addition of 400 nM atto633-*oskar* 3’UTR. Insets are marked with yellow boxes. Naked RNA strands, more prominently visible in condensate periphery in 30 min aged condensates, are marked with black arrowheads. Also refer to movies S11 and S12 for full tomograms. (C) Condensates were assembled with 100 nM *oskar* 3’UTR-atto633 (blue) and EGFP-tagged Bruno or Hrp48 (green). 10 µM RFP-PTB (red) or TMR (tetramethyl rhodamine)-BSA (red) was added after 30 min of incubation. Single confocal slices shown. (D) 2 µM RFP-PTB was added to 30 min aged Bruno-*oskar* 3’UTR condensates and cryo-ET was performed on the condensates. Inset is marked with yellow box. Protein clusters are marked with arrowheads. For full tomogram volumes refer to Movie S13. See also Figure S4.

*oskar* granules *in vivo* show a distribution of sizes with an average diameter of 400 ± 129 nm (Figure 1A). Quantifications of STED data revealed that increase in granule volume correlated with increase in RNA signal. However, there was no net increase in the concentration of RNA with increase in granule volume (Figure S4E). This implies that once hardened, the granules cannot incorporate more *oskar* RNA, highlighting the potential importance of the liquid phase for RNA incorporation during granule assembly.

The observed non-dynamic nature of the condensates both *in vitro* and *in vivo* following ageing raises questions about incorporation of other proteins known to associate with *oskar en route* to the posterior pole, such as PTB (Figure 2A). To test if the solid condensates could incorporate proteins, we added mRFP-PTB to 30 min aged condensates of either Bruno or Hrp48 assembled with *oskar* 3’UTR. Although mRFP-PTB did not phase separate with *oskar* 3’UTR on its own (Figure 2C), it selectively partitioned into the condensates (Figure 3C). This enrichment was not only protein specific (mRFP-PTB vs. TMR-BSA), but also RNA dependent, as Bruno or Hrp48-only condensates did not concentrate PTB (Figure 3C). Cryo-ET revealed that PTB-enriched condensates were amorphous and did not manifest a structural change (Figure 3D, Movie S13).

### Bruno PrLD plays a pivotal role in *oskar* granule assembly

Our *in vitro* reconstitutions show that two *bona fide* granule proteins Bruno and Hrp48 phase separate with *oskar* 3’UTR into liquid-like condensates that rapidly harden into a solid state. PTB, on the other hand, did not form condensates by itself, but only partitioned into preformed *oskar* 3’UTR containing condensates. Additionally, *ptb*-RNAi in the germline had no visible effect on *oskar* granules (Figure S5A-C). This led us to investigate how the intrinsically phase separating Bruno and Hrp48 affect condensation and material properties of the granules *in vivo*. The early association and enrichment of Bruno on *oskar* in nurse cells and its role in higher order particle formation indicate that Bruno may have a central role in granule assembly. However, manipulating levels of Bruno in the germline is detrimental for oogenesis, preventing us from analyzing the effect of Bruno depletion on *oskar* granule assembly (Filardo and Ephrussi, 2003; Webster et al., 1997).

Analysis of the primary sequence revealed that the N-terminal PrLD (Figure 2B) is highly conserved among Drosophilids (Figure S6A). A possible role of the N-terminal in Bruno dimerization was reported (Kim et al., 2015). EGFP-tagged full length Bruno (Bruno FL-EGFP) assembled into distinct granules in *Drosophila* Schneider cells, unlike the N-terminal truncated (Bruno ΔN-EGFP) version (Figure S6B). In contrast, both FL and ΔN proteins phase separated *in vitro* (Figure S6C). To unambiguously address the role of Bruno PrLD *in vivo*, where *oskar* RNA is expressed, we generated transgenic flies expressing Bruno FL-EGFP and Bruno ΔN-EGFP in a tissue-specific manner (Figure 4A). The transgenes were inserted by site-specific integration at an intergenic locus to avoid variability in gene expression owing to chromosomal context. Over-expression of Bruno ΔN-EGFP was toxic to the germline and egg chambers degenerated at early stages (Figure S7A). Therefore, expression of both transgenes was carried out in an endogenous Bruno deficient background (*aret*PA62/*aret*^CRISPR null^). The FL protein rescued oogenesis, while expression of ΔN resulted in only partial rescue of oogenesis, with egg chambers degenerating beyond stage 9 (Figure S7B). Notably, the FL protein formed distinct granules some which co-localized with *oskar* in the nurse cell cytoplasm, with enrichment on track-like structures (Figure 4B and Movie S14). Bruno ΔN-EGFP was largely diffuse, suggesting a role of the N-terminal PrLD in granule formation (Figure 4A-B). Unlike Bruno FL-EGFP, expression of Bruno ΔN-EGFP led to complete failure in *oskar* localization at the posterior pole in stage 9 (Figure 4A). Localization of two other maternal RNAs, *gurken* and *bicoid* was unaffected, confirming the effect was specific for *oskar* mRNA (Figure S8A). Strikingly, *oskar* granules lost their punctate appearance in the case of ΔN, and instead appeared diffuse, indicating an impairment of granule formation (Figure 4B). Quantification revealed a 2-fold reduction in *oskar* enrichment in granules upon ΔN expression compared to FL Bruno (Figure S8B).

**Figure 4.**
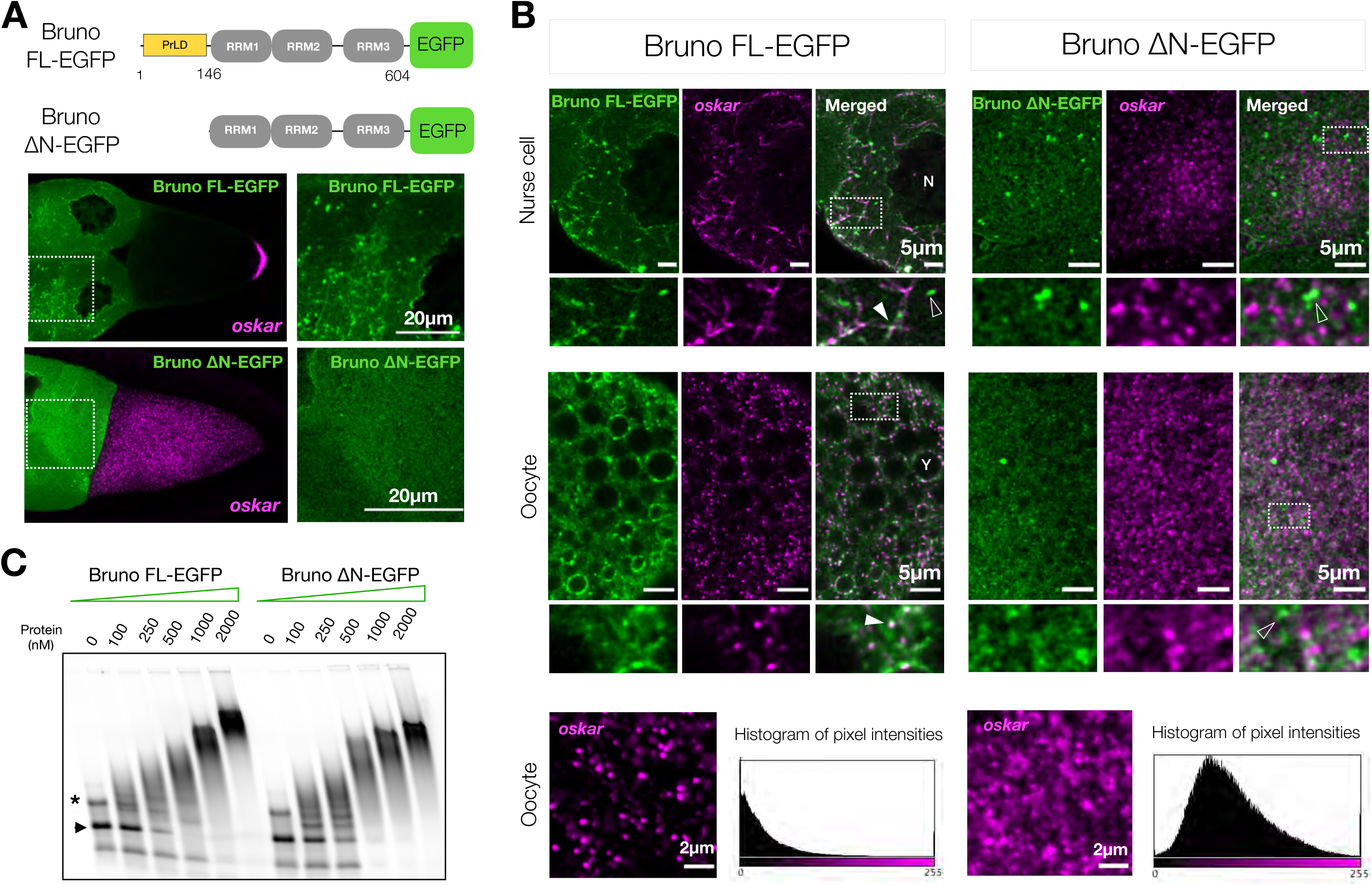
Bruno is essential for *oskar* granule assembly. (A) Schematic representation of Bruno constructs used for transgenesis. Bruno FL-EGFP forms granules in the egg chambers, unlike ΔN which does not. smFISH for *oskar* RNA (magenta) indicates posterior enrichment of *oskar* in stage 9 egg chambers expressing FL Bruno (green), but not ΔN. Boxed regions are enlarged on the right. (B) Single plane confocal images of egg chambers expressing Bruno FL- or ΔN-EGFP (green) and *oskar* (magenta). Insets are marked with a white dotted box. Bold white arrows mark colocalization of protein with *oskar*; empty white arrows indicate protein puncta not associated with *oskar*; N (nurse cell nucleus) and Y (yolk granule). Enlarged view of *oskar* granules (magenta) in ooplasm. Images are acquired with independent microscope settings. A histogram of pixel intensities of the two images confirms the significant loss of granule formation and diffuse *oskar* RNA signal in Bruno ΔN-EGFP. (C) EMSA of fluorescently labelled *oskar* 3’UTR (50 nM) with increasing concentrations of recombinant EGFP-tagged Bruno FL and ΔN suggests that both the full length and truncated protein bind *oskar* 3’UTR with comparable affinity. Arrowhead: *oskar* 3’UTR and *: dimeric form of the 3’UTR. See also figures S5-S8.

We asked whether ΔN Bruno fails to bind *oskar* mRNA and therefore failed to localize with *oskar*. EMSA confirmed that recombinant ΔN, which retains all 3 RRMs of Bruno, is capable of binding *oskar* 3’UTR and forming higher-order oligomers (Figure 4C). This implies that ΔN likely binds *oskar in vivo* but fails to phase separate and form granules. Moreover, Oskar protein was not detected in ΔN egg chambers, unlike the FL (Figure S8C-D). Taken together, our *in vivo* experiments suggest that the scaffold protein Bruno, in particular its PrLD, plays a dominant role in *oskar* granule assembly. *In vivo,* granule assembly precedes hardening, therefore the role of Bruno in hardening could not be addressed in the germline.

### PrLD of Hrp48 is crucial for *oskar* localization and translation

Loss of Hrp48 due to P-element insertion in the gene has been reported to disrupt *oskar* localization (Yano et al., 2004). We carried out a conditional knock down of Hrp48 in the germline by RNAi. Unlike Bruno RNAi, Hrp48 knock down in the germline does not cause early oogenesis arrest, allowing us to score the *oskar* phenotype. Enrichment of *oskar* in the oocyte was unaffected in Hrp48 RNAi early-stage egg chambers (Huynh et al., 2004) (data not shown). Mislocalization was detected in stage 9, when *oskar* accumulated as a cloud in the center of the oocyte (Figure 5A and B). In later stages, larger assemblies of 2-4 µm diameter were detected near the posterior pole, possibly arising from coalescence of smaller granules (Figure 5C). Localization of two other maternal RNAs, *gurken* and *bicoid* was unaffected, confirming that cell polarity was not affected upon Hrp48 RNAi (Figure S9A).

**Figure 5.**
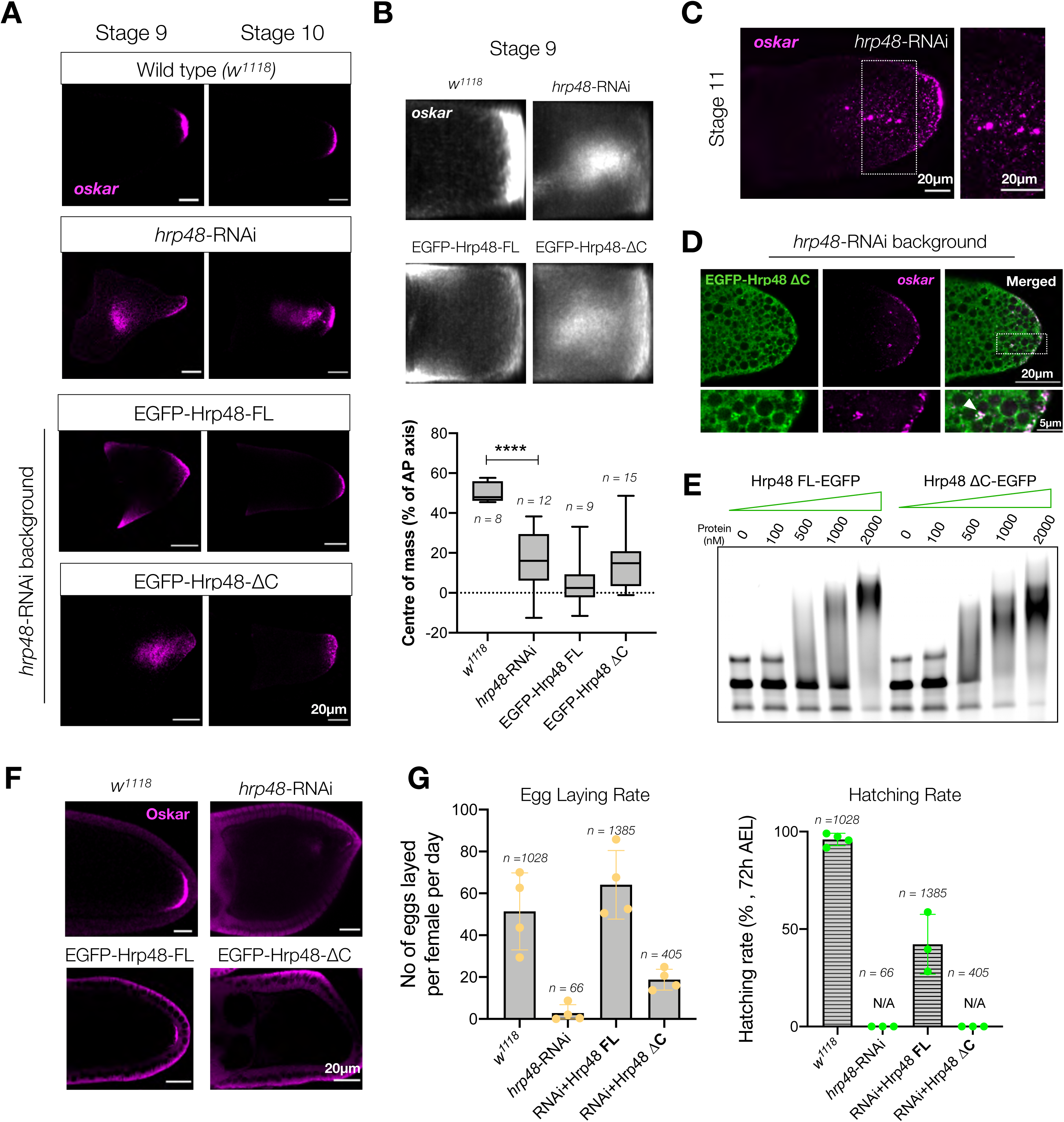
Loss of Hrp48 from the germline impairs *oskar* localization and translation. (A) Detection of *oskar* mRNA (magenta) by smFISH in stages 9 and 10 egg chambers of the indicated genotypes. *w^1118^* is used as wild-type control; RNAi is driven in the germline by *oskar*GAL4 driver; rescue experiments with *oskar*GAL4 driven expression of EGFP-Hrp48 FL and EGFP-Hrp48 ΔC in *hrp48*-RNAi background. (B) Mean *oskar* RNA signal (grayscale) from smFISH data from stage 9 oocytes; anterior to the left. Position of the *oskar* center of mass relative to the geometric center of the oocyte (dotted horizontal line) along the antero-posterior (AP) axis. Error bars represent SD and *n* denotes number of analysed oocytes. Unpaired Student’s t-tests were used for comparisons. Significance level: **** < 0.0001. (C) Clustering of *oskar* mRNA (magenta) into large micron-sized condensates in *hrp48*-RNAi oocytes. Boxed area is enlarged on the right. (D-E) Confocal slice of a cortical plane of stage 10 oocyte show that EGFP-Hrp48 ΔC (green) associates with *oskar* (magenta) granules (white arrow). Boxed area is enlarged below (D). EMSA of fluorescently labelled *oskar* 3’UTR (50 nM) with increasing concentrations of recombinant EGFP-tagged Hrp48 FL and ΔC confirms that ΔC binds *oskar* 3’UTR (E). (F) Immunostaining of Oskar protein (magenta); signal in follicle cells is background from the antibody and is detected in Oskar protein null egg chambers (Figure S8D). (G) Egg laying and hatching rate from flies of the indicated genotypes show that loss of Hrp48 by RNAi impairs egg laying and hatching, which is not rescued by expression of ΔC protein. Error bars represent SD. See also figure S9.

Mutations within the PrLD of Hrp48 were reported to impair *oskar* localization and translation, but not binding to *oskar* mRNA (Huynh et al., 2004). To test the importance of the PrLD, we generated transgenic flies expressing EGFP-tagged full-length or a PrLD-truncated version (ΔC) of Hrp48 (Figure S9B) and scored their ability to rescue the RNAi phenotype. At stage 9, the clustering of *oskar* granules in the center of the oocyte was not rescued by expression of ΔC protein (Figure 5A). Expression of the FL protein rescued the central clustering, but anterior mislocalization was observed possibly due to higher expression levels of Hrp48 FL compared to ΔC (Figure S9B). At stage 10, posterior localization of *oskar* was rescued in FL oocytes, while the ΔC phenotype resembled that of the RNAi, with larger granules dispersed near the posterior pole (Figure 5A and S9C). The PrLD-truncated Hrp48, which retains the two RRMs, localized with *oskar in vivo* (Figure 5D). EMSA confirmed that the ΔC protein binds *oskar* forming higher-order oligomers *in vitro* (Figure 5E). Translation of Oskar protein was impaired in the *hrp48*-RNAi flies and egg laying was drastically reduced (Figure 5F-G). Expression of ΔC failed to rescue translation (Figure 5F), while egg laying was only partially rescued and the eggs failed to hatch (Figure 5G).

Depletion of Hrp48 from the germline or the expression of ΔC version, however, did not abolish *oskar* granule formation, presumably owing to Bruno-driven LLPS. Instead, the granules appeared to coalesce into larger condensates. This suggests that *oskar* granules in Hrp48-depleted/truncated lines have altered physical properties that impair their native function, and imply a role of the Hrp48-PrLD in modulating granule material properties. Similar observation has been made in case of Imp, a conserved component of *Drosophila* neuronal RNP granules (Vijayakumar et al., 2019). Biomolecular condensates exhibit a continuum of material and emergent properties that can be harnessed by the cell to specific needs (Alberti, 2017). Therefore, key questions arise as to the significance of the solid state in the physiological function of *oskar* in the germline.

### Manipulating the material state of *oskar* granules impairs RNP posterior localization

To address the importance of the solidification following LLPS, we sought to drive the material properties of *oskar* granules towards a more liquid state without interfering with the endogenous scaffold proteins and the multitude of other functions they perform in the germline. We opted for tethering of an exogenous low complexity (LC) domain of human FUS, which reversibly phase separates into liquid condensates in living cells and *in vitro* forms highly dynamic liquid droplets that flow, fuse and spontaneously relax into a spherical shape (Burke et al., 2015; Shin et al., 2017). We genetically tethered the FUS LC domain to *oskar* mRNA, using the MCP-MS2 system.

We generated transgenic flies expressing MCP-EGFP-FUS LC and a control line where the FUS LC is replaced by another molecule of EGFP (Figure 6A) to ensure that the expressed proteins are of comparable molecular weight. Expression of these constructs in absence of MS2-tagged *oskar* had no effect on oogenesis and embryogenesis, as the transgenic flies propagate as stable lines (Figure S10A). For genetic tethering, we crossed in an *oskar*6xMS2 transgene (Lin et al., 2008) that rescues oogenesis and embryogenesis in an *oskar* RNA null genetic background. We used an *oskar* RNA null background (*oskA87*/*Df3Rp^XT103^*) in our experiments, as MCP-bound *oskar* transcripts can hitchhike on the endogenous *oskar* RNA (Hachet and Ephrussi, 2004; Jambor et al., 2011).

**Figure 6.**
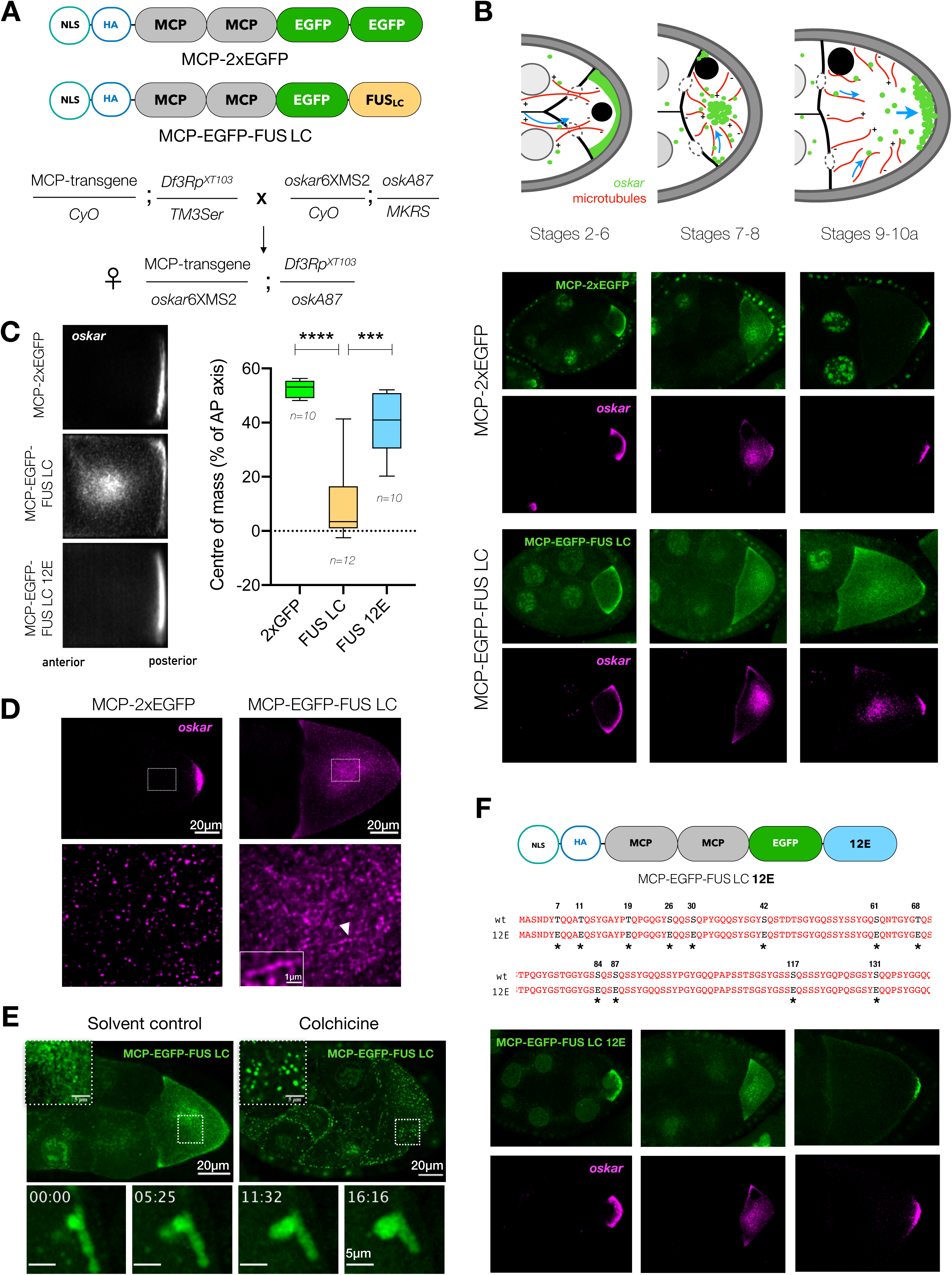
Manipulating the material properties of *oskar* granules affects RNA localization. (A) Constructs used for transgenesis and scheme of the genetic crosses. (B) Cartoon representation of *oskar* RNA localization at distinct stages of oogenesis (adapted from (Cha et al., 2002)). Representative confocal images of *oskar* distribution from early to mid-oogenesis upon 2xEGFP and FUS LC tethering; EGFP (green) and *oskar* (magenta). (C) Quantification of transport defects. Mean *oskar* RNA signal (grayscale) from smFISH data from stage 9 oocytes; anterior to the left. Position of the *oskar* center of mass relative to the geometric center of the oocyte (dotted horizontal line) along the AP axis. Error bars represent SD and *n* denotes number of oocytes analysed. Unpaired Student’s t-tests were used for comparisons. Significance levels: *** < 0.001, **** < 0.0001. (D) Stage 9 egg chambers with *oskar* RNA (magenta) detected by FISH; a central region in the oocyte (dotted white box) is enlarged below. White arrowhead marks track-like structure with *oskar* granules. (E) Depolymerizing microtubules with colchicine in isolated ovaries results in assembly of large spherical granules in FUS LC tethered condition (upper panel). Lower panel depicts liquid-like behavior of FUS LC-*oskar* granules upon colchicine treatment (Movie S16). (F) Cartoon representation of the rescue construct MCP-EGFP-FUS LC 12E with the 12 mutated sites marked with *. Representative images of early to mid-oogenesis stages in 12E tethered condition; MCP-EGFP-FUS LC 12E (green) and *oskar* (magenta). See also figures S10 and S11.

FUS LC-tethered *oskar* granules were transported from the nurse cells to the oocyte and rescued oogenesis similar to the 2xEGFP tethered control (Figure 6B). Therefore, FUS LC-tethering did not impair granule association with microtubules or the dynein machinery. At stages 7-8, the oocyte microtubule network reorganizes and *oskar* granules are transported by kinesin, initially away from the cortex to the interior, and eventually to the posterior pole at stage 9 (Cha et al., 2002). Movement of *oskar* granules towards the interior of the developing oocyte was indistinguishable between 2xEGFP or FUS LC tethering (Figure 6B and S10B). However, in stage 9 egg chambers, while the 2xEGFP-tethered *oskar* localized to the posterior pole, the FUS LC-tethered *oskar* was severely mislocalized: *oskar* appeared as a cloud in the center of the oocyte (Figure 6B and C). This phenotype was similar to the mislocalization observed upon Hrp48 knockdown/truncation (Figure 5A-B). Live imaging revealed directed runs of FUS LC-tethered *oskar* granules, indicating that microtubule association per se is not affected upon FUS LC tethering (Figure S10C; Movie S15). Therefore, mislocalization cannot be attributed to lack of kinesin association or loss of microtubule association.

Careful examination of the cloud-like mass of *oskar* upon FUS LC tethering showed occasional presence of larger granules and track-like segments with granules aligned like beads on a string (Figure 6D). We hypothesized that microtubule-directed transport of the granules prevents them from fusing into larger condensates and accumulating at ectopic locations. Depolymerization of microtubules *ex vivo* by colchicine resulted in collapse of the diffraction-limited granules into large structures both in the oocyte and nurse cells. The granules were found to fuse and relax like liquids, and to wet membrane surfaces demonstrating the liquid properties of FUS LC-tethered *oskar* granules (Figure 6E, S10D, Movie S16). We further exposed the colchicine treated egg chambers to 1,6-hexanediol which partially dissolved the large granules (Figure S10E). Existence of the smaller granules indicated that the original solid phase persisted.

Phosphorylated and a phosphomimetic version of FUS LC have been shown to ablate its phase separation potential *in vivo* and *in vitro* (Monahan et al., 2017; Rhoads et al., 2018; Shorter, 2017). We reasoned that if the *oskar* localization defects arise from the induced liquid state of the granules, tethering a phase separation deficient form of FUS LC should restore posterior localization. We generated transgenic flies expressing the phosphomimetic version of the LC (MCP-EGFP-FUS LC 12E). In contrast to wild type FUS LC, the phase separation-compromised version induced no aberrant central accumulation of *oskar* in stage 9 oocytes (Figure 6C, F and S10F).

The FUS LC-induced localization defects were also absent when an endogenous copy of the *oskar* gene was provided in addition to the *oskar*6xMS2 transgene. By virtue of 3’UTR mediated hitchhiking, the *oskar*6XMS2 mRNA and the endogenous *oskar* transcripts co-package into same granules (Figure S11A), diluting the relative protein-to-mRNA ratio and resulting in reduced FUS LC phase separation (Figure S11B).

### The solid state is important for *oskar* mRNA translation

*oskar* mRNA is translationally repressed during transport. De-repression upon localization results in Oskar protein production at the posterior pole. Whereas a posterior crescent of *oskar* mRNA is present in the control 2xEGFP, FUS LC-*oskar* granules fail to form a crescent-shaped structure (Figure 7A). Instead, local concentration of FUS LC above the critical concentration for condensation leads to formation of spherical, micron-sized condensates as observed upon loss of Hrp48 (Figure 5C). The large condensates were loosely anchored and subsequently detached into the ooplasm (Figure 7A-B and Movie S17).

**Figure 7.**
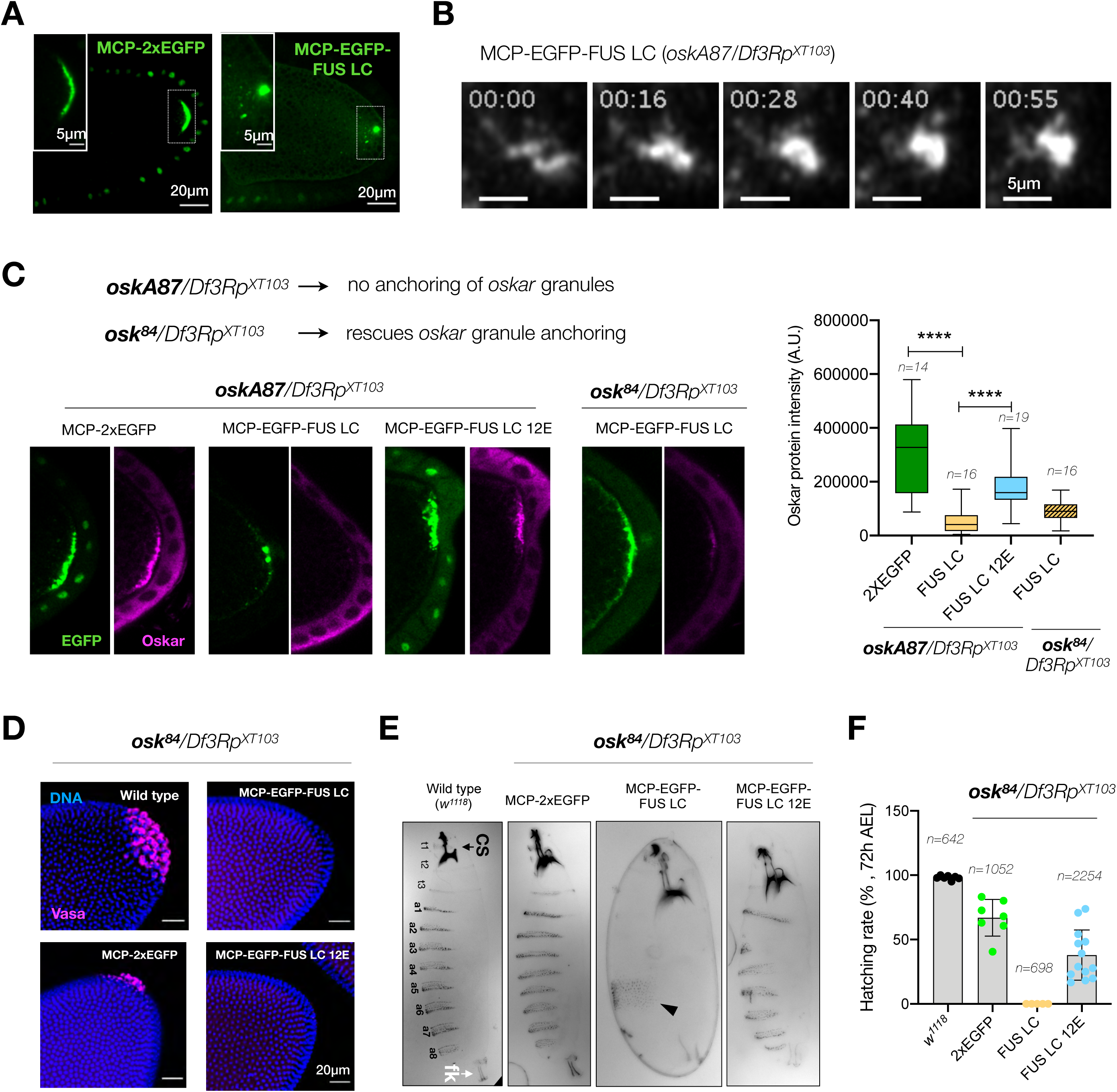
Manipulating the material properties of *oskar* granules interferes with *oskar* anchoring and translation, and impairs embryonic development. (A) Representative stage 10 egg chambers expressing MCP-2xEGFP or MCP-EGFP-FUS LC in an *oskar* RNA null background (*oskA87*/*Df3Rp^XT103^*) show loss of granule anchoring. (B) Movie snapshots of fusion of *oskar* granules tethered with MCP-EGFP-FUS LC (grayscale) in an *oskar* RNA null background (*oskA87*/*Df3Rp^XT103^*) near the posterior cortex. Also refer to Movie S17. (C) Immunostaining for Oskar protein in *oskar* RNA null (*oskA87*/*Df3Rp^XT103^*) egg chambers. The genotype of the fourth panel is *osk^84^*/*Df3Rp^XT103^* where the anchoring function is provided *in trans*; EGFP (green) and Oskar protein (magenta). Refer to S12F for all conditions in the *osk^84^*/*Df3Rp^XT103^* background. All images shown (and used for quantification) were acquired using identical microscope settings and representations are contrast-matched. Quantification of the Oskar signal intensity at the posterior pole of several egg chambers. Error bars represent SD and *n* denotes number of analysed oocytes. Unpaired Student’s t-tests were used for comparisons. Significance level: **** < 0.0001. (D) Representative images of the posterior of embryos of the indicted genotypes at nuclear cycle 14. Pole cells are identified by Vasa immunostaining (magenta) and nuclei are stained with DAPI (blue). (E) Cuticles of embryos of the indicated genotypes; anterior at the top, ventral to the left. t1-t3: thoracic segments; a1-a8: abdominal segments; cs: cephalo-pharyngeal skeleton (black arrow); fk: Filzkörper material (white arrow); black arrowhead: patchy band of denticles. (F) Quantification of the hatching rates of eggs from females expressing the indicated transgene in *osk^84^*/*Df3Rp^XT103^* background. Error bars represent SD and *n* denotes the number of eggs scored per genotype. See also figures S12 and S13.

Similar to the *hrp48*-RNAi phenotype, upon FUS LC-tethering Oskar protein was not detected at the posterior pole (Figure 7C). In contrast to FUS LC, the 12E construct only partially impaired Oskar translation (Figure 7C). Western blotting confirmed reduction in levels of both the Long and Short Oskar isoforms (Figure S12A-B). Short Oskar induces assembly of the pole plasm (Ephrussi and Lehmann, 1992) and the Long isoform organizes the posterior cortex of the oocyte and anchors the pole plasm (Vanzo et al., 2007; Vanzo and Ephrussi, 2002)(Figure S12C). This made us question whether the observed detachment of FUS LC-tethered *oskar* granules is a consequence of loss of anchoring due to reduced Oskar protein levels, resulting in further reduction of *oskar* mRNA translation.

The N-terminus of Oskar is sufficient for anchoring (Hurd et al., 2016; Vanzo and Ephrussi, 2002). To uncouple the interdependency between anchoring and translation, we provided the anchoring function *in trans* using an *oskar* nonsense mutant allele, *osk^84^*, which encodes a 254 residue long peptide from the N-terminus (Figure S12A). The *osk^84^* nonsense mutant completely rescued the anchoring function at stage 9-10, with minor delocalization only in late stages (Vanzo and Ephrussi, 2002)(Figure S12D). The presence of this additional copy of *oskar* also suppressed the transport defects, as observed with a wild type *oskar* allele (Figure S11B). The pole detachment phenotype observed in 90% of stage 10b egg chambers upon FUS-tethering was rescued when anchoring was provided, and the granules did not collapse into large condensates (Figure S12E). However, translation into Oskar protein was still compromised with FUS LC tethering, even when anchoring was rescued with *osk^84^* (Figure 7C and S12F). Thus, the liquid-like phase, rather than defective anchoring, is responsible for the observed translational shutdown.

### Altered material state of *oskar* granules is detrimental to embryonic development

A localized source of Oskar is crucial for germline development and antero-posterior patterning of the embryo (Lehmann and Nusslein-Volhard, 1986). Whereas a small amount of Oskar is sufficient for patterning the abdomen, a high local concentration of the protein is required to induce germ cell formation (Ephrussi and Lehmann, 1992; Smith et al., 1992). To investigate the effect of changed material properties of *oskar* granules on germline formation, we assessed pole cell formation in embryos at the blastoderm stage. While wild type (*w^1118^*) embryos had an average of 25-30 pole cells, 2xEGFP tethering to *oskar*6xMS2 RNA in an *oskar* RNA null background resulted in 10-12 pole cells. FUS LC tethering led to complete absence of pole cells, whether or not anchoring was provided *in trans* by the *osk^84^* allele. Though Oskar was detected in FUS LC 12E expressing oocytes, the amount of protein was not sufficient to induce pole cell formation in both *oskA87* and *osk^84^* genetic backgrounds (Figure 7D and S13A).

We additionally analyzed antero-posterior patterning of the embryos. Tethering of 2xEGFP resulted in formation of all 8 abdominal segments (a1-a8), as in wild type embryos (Figure 7E and S13B), and no head defects were observed. This indicates that the pole plasm was appropriately localized and its abdomen-inducing activity was comparable to wild type. Upon FUS LC tethering, the embryos displayed a loss of abdominal segmentation, as reported for strong loss-of-function *oskar* alleles (Lehmann and Nusslein-Volhard, 1986). Rescue of anchoring by complementation with the *osk^84^* allele did not rescue abdominal segmentation upon FUS LC tethering. Quantification of larval hatching rates revealed that 66% of the 2xEGFP embryos hatched, while FUS LC embryos did not hatch at all, whether in an *oskA87/Df3Rp^XT103^* or *osk^84^/Df3Rp^XT103^* background (Figure 7F and S13C). In the *osk^84^/Df3Rp^XT103^* background, expression of FUS LC 12E resulted in a remarkable 44% of hatching, with embryos exhibiting 4-6 abdominal segments. This underlines how modulation of physical state of *oskar* granules towards a more liquid phase impacts the development of the future embryo.

## Discussion

Our study of *oskar* transport granules in the *Drosophila* female germline highlights the significance of regulating material properties of RNP granules for their physiological functions. We identify scaffold proteins, implicated in translation regulation, that *in vitro* play a major function in driving granule formation via LLPS and liquid-to-solid phase transition. *In vivo oskar* granules exhibit spherical morphology with solid-like material properties. It is likely that the initial liquid-like phase is short-lived and could not be detected. To investigate the significance of condensate solidification, we perturb the solid state of the granules and show the importance of the solid material properties in granule localization and translation. Our findings not only elucidate key principles of *oskar* granule assembly, but also reflect how cells assemble a stable and apparently non-dynamic condensate to regulate localization and expression of a maternal RNA that is a master regulator of germline formation.

Functional biomolecular condensates exhibit a spectrum of material physical states. While the nucleolus, stress granules and P-granules are liquid-like, Balbiani bodies and centrosomal PCM are more solid-like (Audas et al., 2016; Boke et al., 2016; Brangwynne et al., 2009; Lafontaine et al., 2020; Patel et al., 2015; Schmidt and Görlich, 2015; Woodruff et al., 2017; Woodruff et al., 2018). Our *in vitro* reconstitutions show that *oskar* minimal condensates assembled with core proteins rapidly mature into a non-dynamic state with respect to fusion or molecular exchange. For *oskar* granules, the liquid state appears to be important for RNA incorporation, as solidification precludes RNA entry into the condensates. Solid *oskar* granules *in vivo* exhibit a range of size distributions. As condensates lack deterministic stoichiometry of the constituent macromolecules, they represent assemblies of variable sizes. *oskar* granules constitute transport cargoes that travel distances up to 100 µm to attain localization. Artificially inducing a liquid-like state drastically compromised the localization efficiency of *oskar* (Figure 6C). Therefore, it is plausible that hardening through non-covalent cross-linking of scaffold proteins and RNA confers mechanical stability that fortifies *oskar* granules with properties that support long distance transport (Figure 2E).

Within the domain of solid-like condensates there exist two classes. On one hand, Balbiani bodies and amyloid-bodies are implicated in dormancy and shut down of chemical reactions by physical sequestration of the components (Audas et al., 2016). On the other hand, the mature PCM, although constituted of a non-dynamic scaffold, allows selective partitioning of effector molecules that are needed for its function. Woodruff et al. (2018) describe such selectively permeable condensates as bio-reactive gels. *oskar* granules belong to the latter class. Imaging of the condensates at molecular resolution with cryo-ET revealed an amorphous appearance, confirming that the initial liquid phase hardens into a glassy solid. Although rapidly developing into non-dynamic condensates, the specific recruitment of *oskar* binding protein PTB into the aged condensates suggests that they are capable of selectively enriching client RNA-binding proteins (Figure 3C). The fact that PTB enriches inside *in vitro* condensates in an *oskar* RNA-dependent manner explains how other crucial components, like Staufen, which do not associate in the nurse cells can be co-detected with *oskar* in the ooplasm (Little et al., 2015). Thus, a first layer of selectivity can arise from relative RNA binding affinities of proteins, as well as the availability of binding sites. Another layer of selectivity presumably arises from the porosity of the condensate. Our experiments show that a 100 kDa protein like RFP-PTB can partition into the condensates, while the ∼330 kDa *oskar* 3’UTR itself is excluded. Therefore, it is expected that small globular proteins can enrich in an RNA-dependent manner, but megadalton complexes like ribosomes are excluded, ensuring translation repression.

Stereospecific molecular interactions seed RNA-protein complexes/oligomers (Little et al., 2015), which by virtue of further multivalent interactions and molecular crowding *in vivo* form mesoscopic assemblies with emergent properties. Bruno and Hrp48 are classic examples of RBPs with a modular architecture: a disordered prion-like domain and structured RNA binding domains. On one hand, RRM-driven sequence-specific binding to *oskar* mRNA ensures selection of specific mRNA, while PrLD-driven phase separation promotes granule assembly via self-association as well as multivalent interactions with other proteins bound to the mRNA. We verified that in absence of their PrLDs, both Bruno and Hrp48 can bind *oskar* mRNA and promote formation of higher-order oligomers (Figure 4C, 5E), but functions pertaining to *in vivo* granule formation and granule material properties are affected. *oskar* mRNA has been shown to dimerize by kissing-loop interactions between stem-loop structures in its 3’UTR (Hachet and Ephrussi, 2004; Jambor et al., 2011). This raises the question whether the mRNA has an architectural role in granule assembly as has been proposed for long non-coding RNAs (lncRNAs) and a few other classes (Yamazaki et al., 2019) (Jain and Vale, 2017). It is plausible that stem-loop mediated RNA-RNA dimerization promotes granule nucleation as has been proposed for *Drosophila bicoid* and HIV genomic RNA (Ferrandon et al., 1997; Paillart et al., 1996; Van Treeck and Parker, 2018; Van Treeck et al., 2018). However, our *in vitro* reconstitutions indicate that self-assembling scaffold proteins like Bruno promote condensation of *oskar* 3’UTR under conditions in which *oskar* 3’UTR alone does not condense into visible assemblies.

In a large polarized cell like the developing oocyte, dynamic microtubule-network organization is essential for transport of maternal RNAs, proteins and organelles to support future embryogenesis. A growing number of examples illustrate the importance of biophysical interactions between condensates and the cytoskeleton. While cytoskeletal fibers can act as platforms that promote condensation by increasing local concentrations (Hernandez-Vega et al., 2017; Wiegand and Hyman, 2020), viscoelastic fibers can restrict condensate dynamics and fusion. Such cytoskeleton-driven spatial segregation of condensates has been observed in the *Xenopus laevis* oocyte nucleus, where a nuclear actin network prevents sinking and fusion of nucleoli, and also recently documented keratinocytes (Feric and Brangwynne, 2013; Quiroz et al., 2020). FUS LC-tethered *oskar* granules appear to exhibit a behavior similar to the latter (Figure 6D). Technical difficulties due to ooplasm viscosity, imaging depth and diffraction-limited granule size prevented us from obtaining higher resolution images of their dynamics. However, depolymerizing the microtubule network by drug treatment confirmed that active transport on microtubule tracks restricts the coalescence of the small granules into larger ones (Figure 6E).

FUS LC-tethering induced a liquid-like state of *oskar* granules and impaired translation at the posterior pole, leading to a cascade of developmental defects. Previously, phase separation of low complexity domain of Fragile X Mental Retardation Protein (FMRP_LC_) has been correlated with translation inhibition in an *in vitro* translation set up (Tsang et al., 2019). It is enigmatic how a change in the physical state of granules can have a dramatic effect on translation control. Due to the lack of knowledge on derepression mechanisms of *oskar* mRNA, we could not identify the cause of the observed impairment of translation. A constitutively active Protein Kinase A (PKA) mutant has been shown to induce ectopic translation of Oskar protein (Yoshida et al., 2004). However, targets of PKA have not been identified. The N-terminal domain of Bruno has been reported to be phosphorylated by PKA *in vitro* (Kim et al., 2015). Therefore, a PKA-driven phosphorylation of scaffold proteins might trigger modulation of condensate architecture and initiate *oskar* translation. An altered physical state may interfere with such a mechanism, either by excluding the kinase or other putative de-repressors. Our attempts to dissolve the colchicine-induced large FUS LC granules by 1,6-hexanediol resulted in partial dissolution (Figure S10E) indicating a possibility of internal compartmentalization. At the posterior pole, where FUS LC concentration surpasses the critical concentration for condensation, micron-size condensates appear. It is possible that FUS LC forms a liquid phase that encapsulates the core RNPs and reduces accessibility of translation factors/derepressors. The mechanism of translational depression is unclear and understudied due to the highly complex relationship between translation, endocytosis, anchoring and pole plasm assembly. It therefore remains an open question how the liquid-like state impaired translation mechanistically.

The persistent presence of *oskar* RNA is toxic to pole cells and it is actively segregated from the germ granules in the posterior domain of the oocyte and embryo (Eichler et al., 2020; Little et al., 2015). A liquid state of the *oskar* granules is therefore a potential threat to this segregation, as co-condensation of *oskar* granules and germ granules would result in co-packaging of *oskar* RNA and other pole cell-destined maternal transcripts. Transition of *oskar* granules into a solid therefore provides a mechanism to ensure proper localization and selective partitioning that are key to the development of oocyte and embryo.

## Acknowledgements

We acknowledge the EMBL Advanced Light Microscopy Facility (ALMF), especially Marko Lampe for help with STED imaging and Aliaksandr Halavatyi for FRAP analysis plugin. We thank the EMBL Electron Microscopy Core Facility (EMCF) and cryo-EM Platform for their support. We acknowledge Alessandra Reversi (*Drosophila* Injection Service, EMBL) for fly injections and the EMBL Protein Expression and Purification Core Facility for providing cloning and expression vectors and TEV protease. We thank Xiaojie Zhang and Anthony Hyman for the FUS plasmid and purified hFUS protein, Bernhard Hampoelz and Martin Beck for the RFP-Nup107 fly line, Tse-Bin Chou and Elizabeth Gavis the *oskar*6xMS2 fly line, Imre Gaspar and Frank Wippich for the RNA copy number calculation R-script. We thank Simon Alberti, Florence Besse and Jeff Woodruff for critical comments on the manuscript. M.B. was supported by a fellowship from the EMBL Interdisciplinary Postdoctoral (EIPOD) Programme under Marie Skłodowska-Curie Actions COFUND (EI3POD). A.E and J.M. acknowledge funding from the EMBL and J.M. from the European Research Council (ERC 3DCellPhase^-^ 760067).

## Author Contributions

All authors conceived the study. M.B. designed and performed the experiments and analyzed the data. Cryo-electron microscopy and analysis was performed by J.M. All authors contributed to the interpretation of the results and wrote the manuscript.

