## Supplementary Material for "Liquid-to-solid phase transition of *oskar* RNP granules is essential for their function in the *Drosophila* germline"

### Supplementary Figures

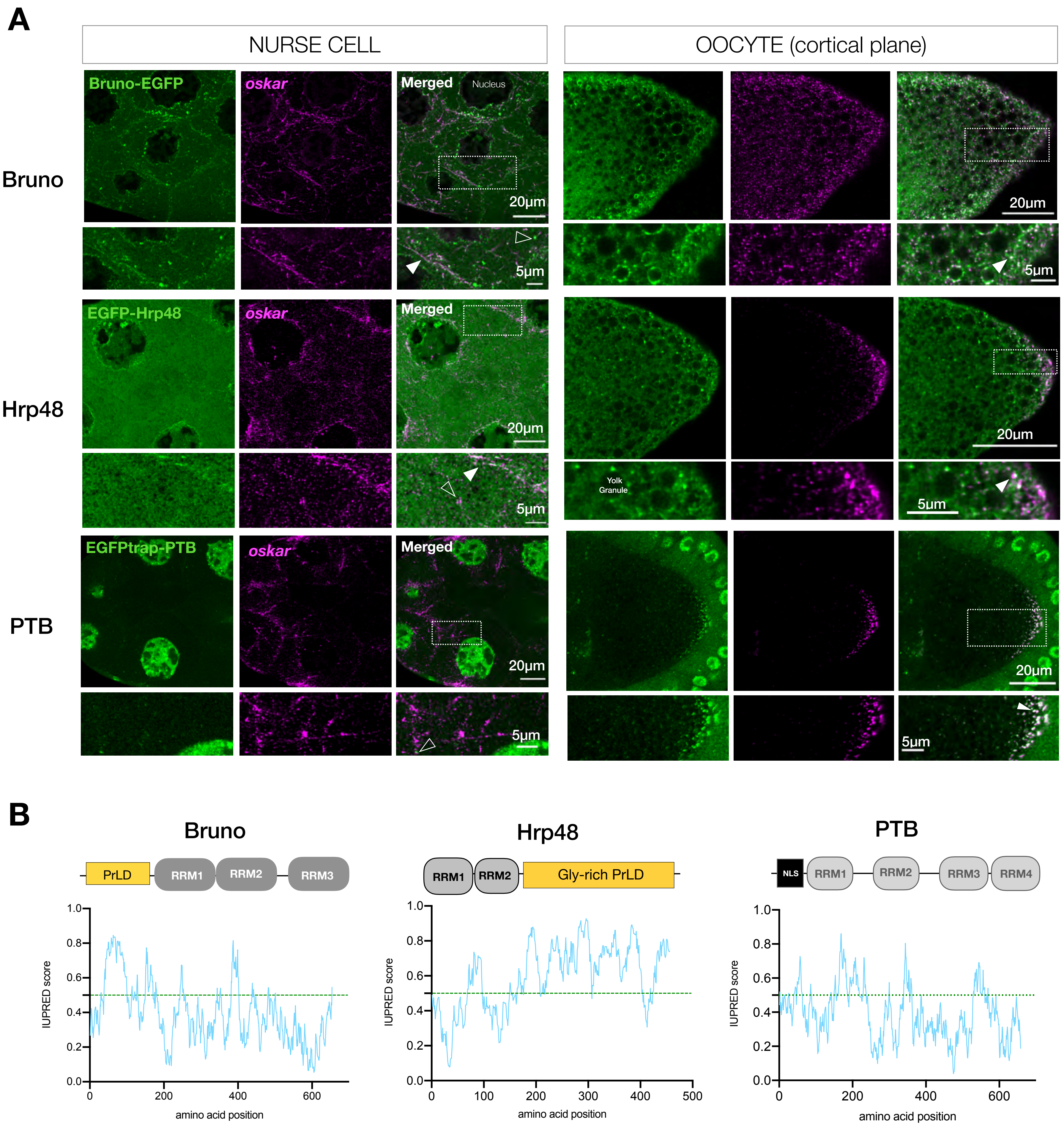

**Figure S1, Related to Figure 2**

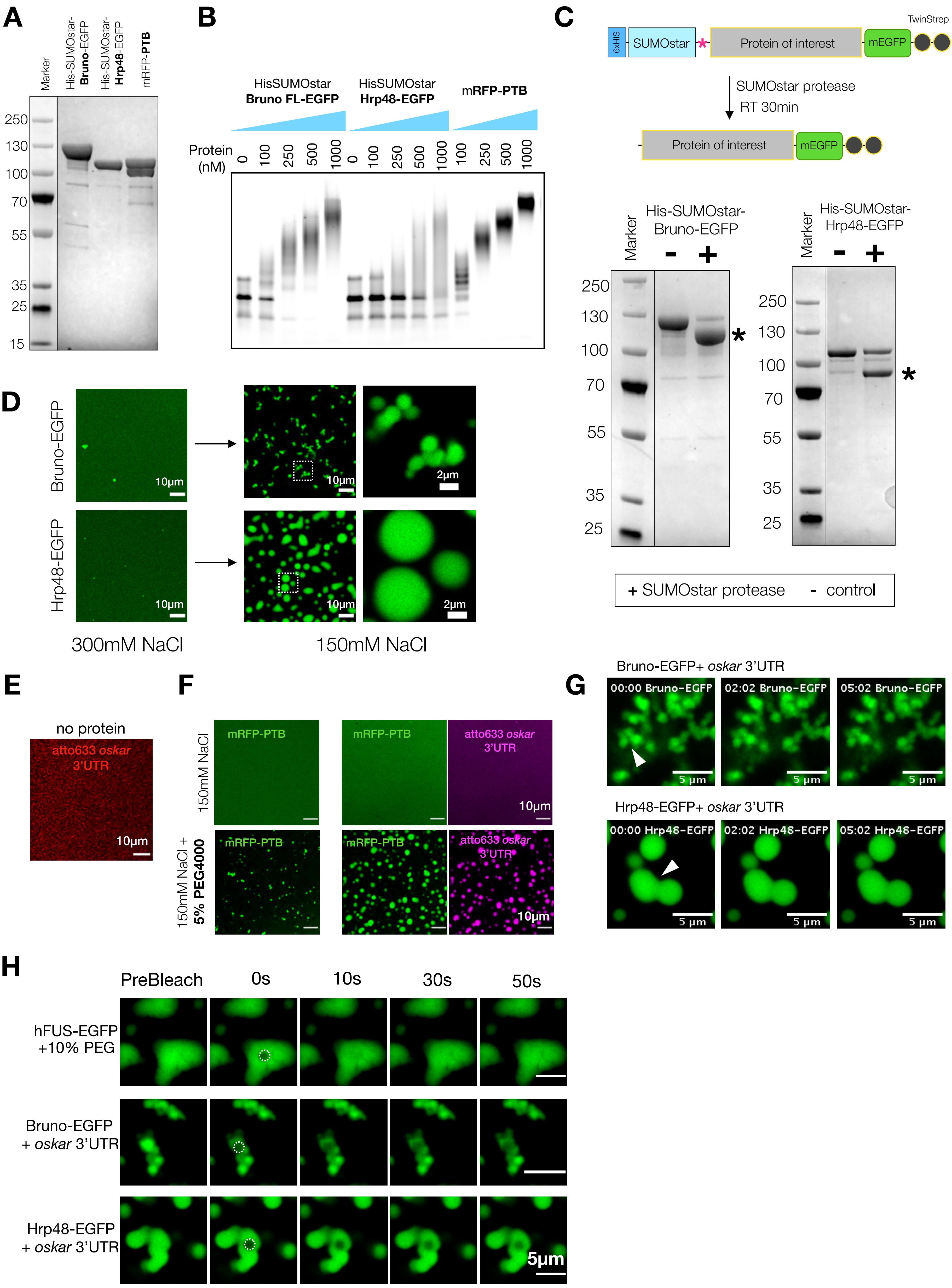

**Figure S2, Related to Figure 2**

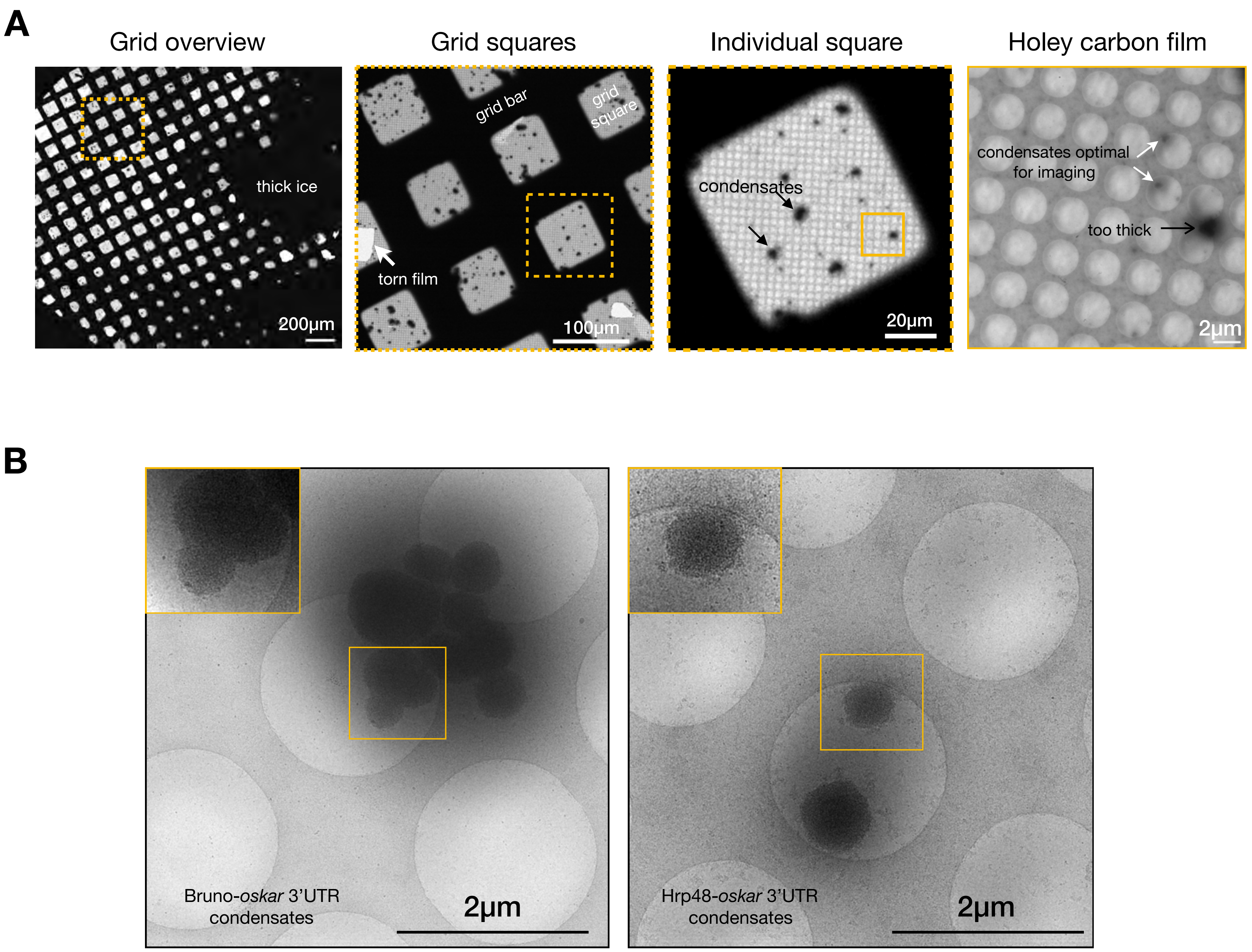

**Figure S3, Related to Figure 2**

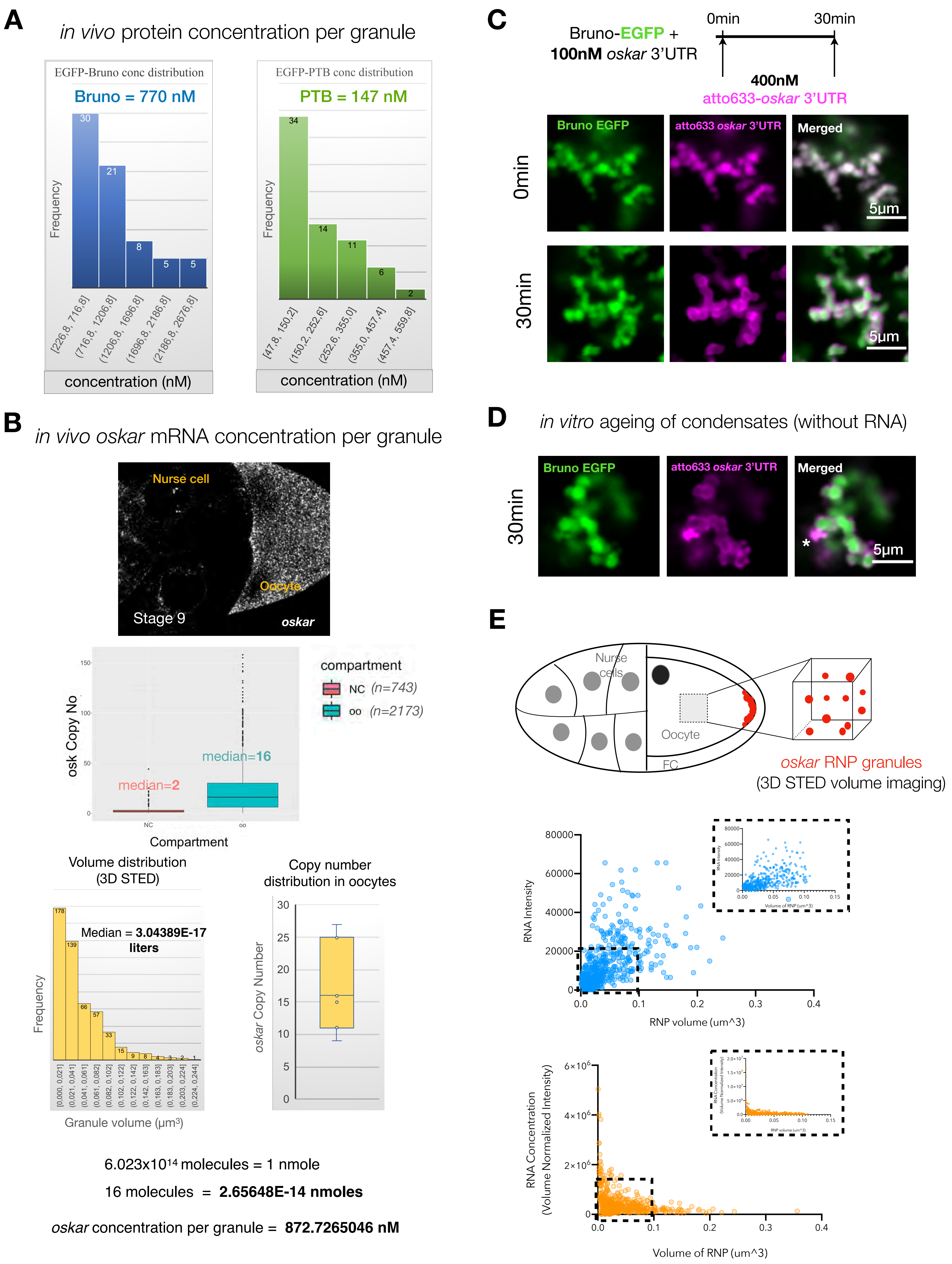

**Figure S4, Related to Figure 3**

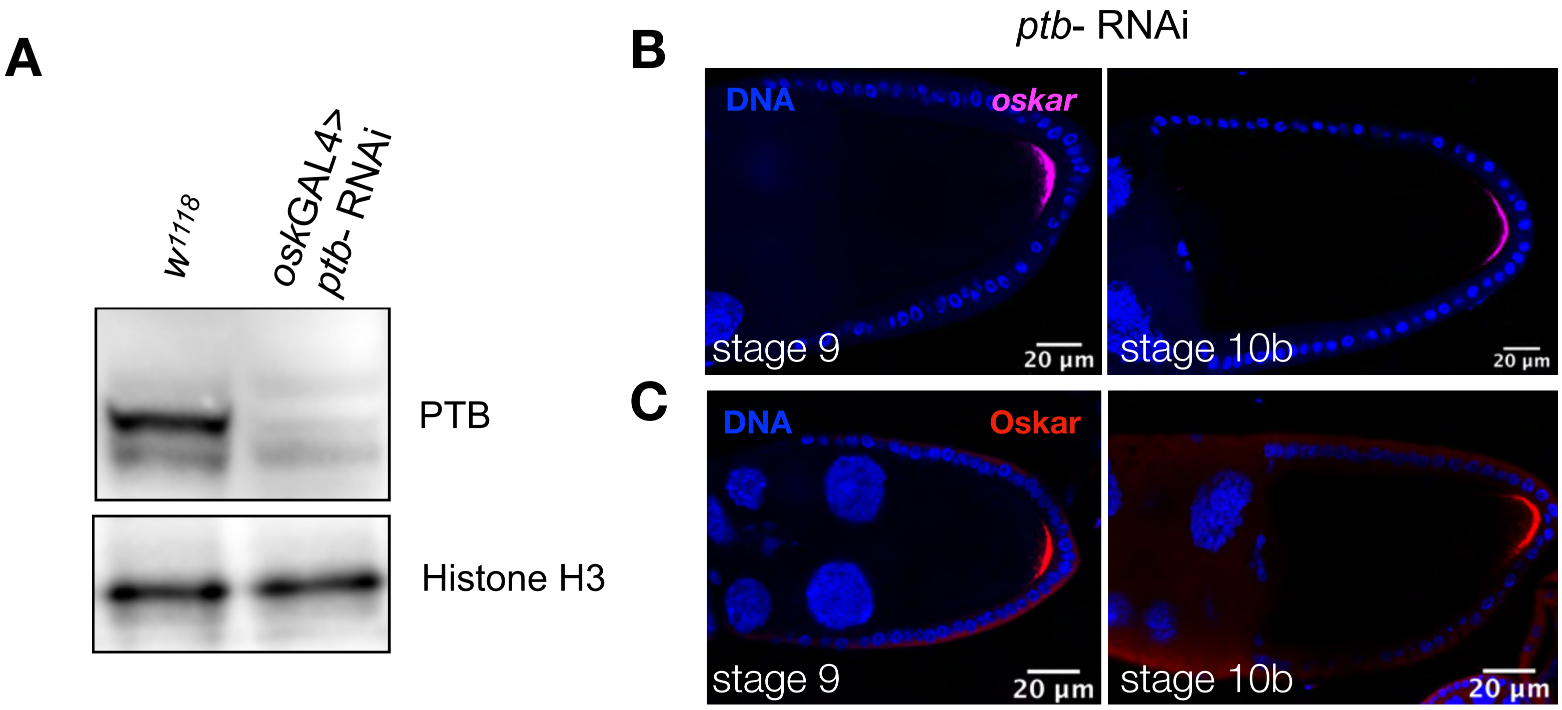

**Figure S5, Related to Figure 4**

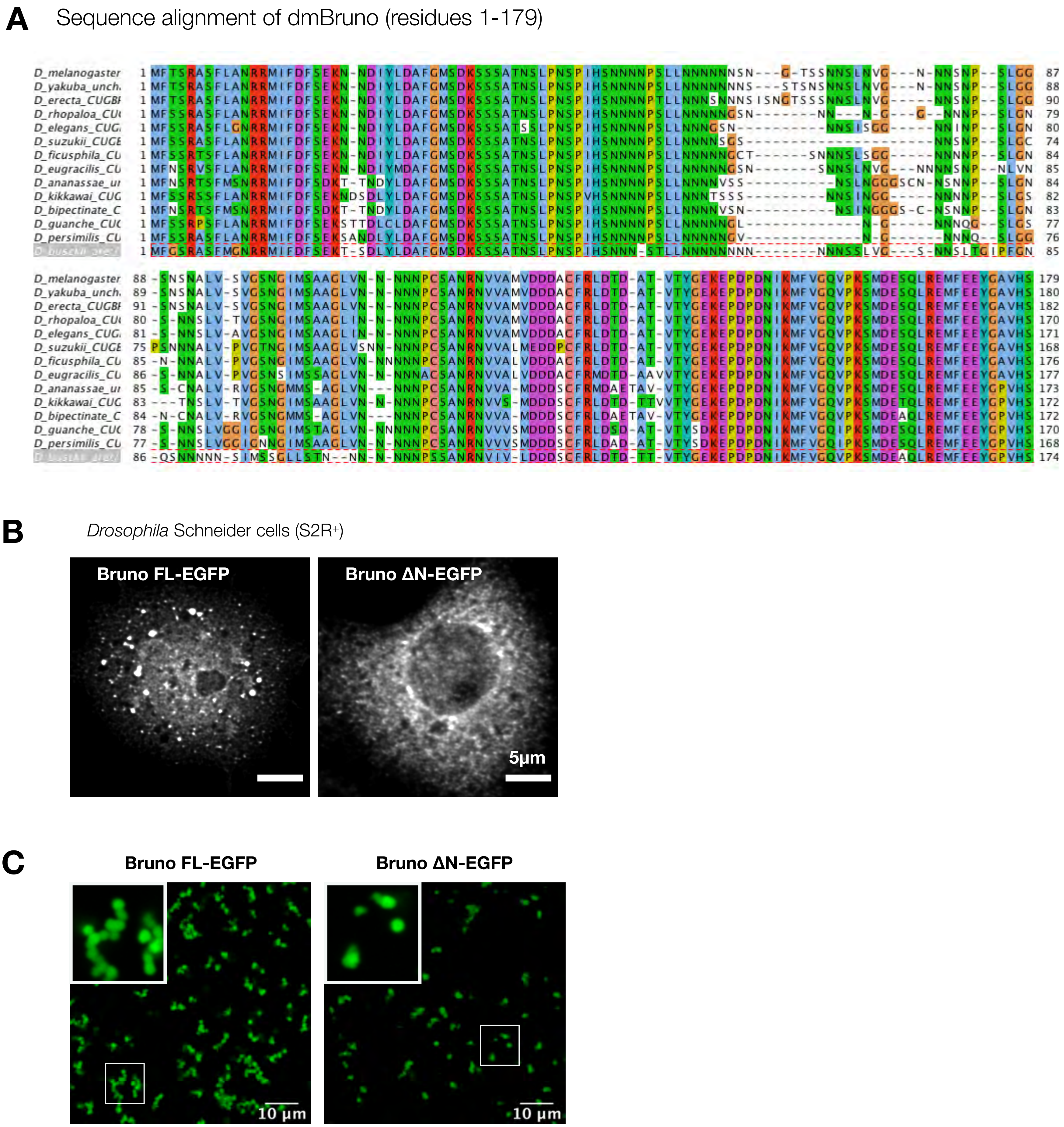

**Figure S6, Related to Figure 4**

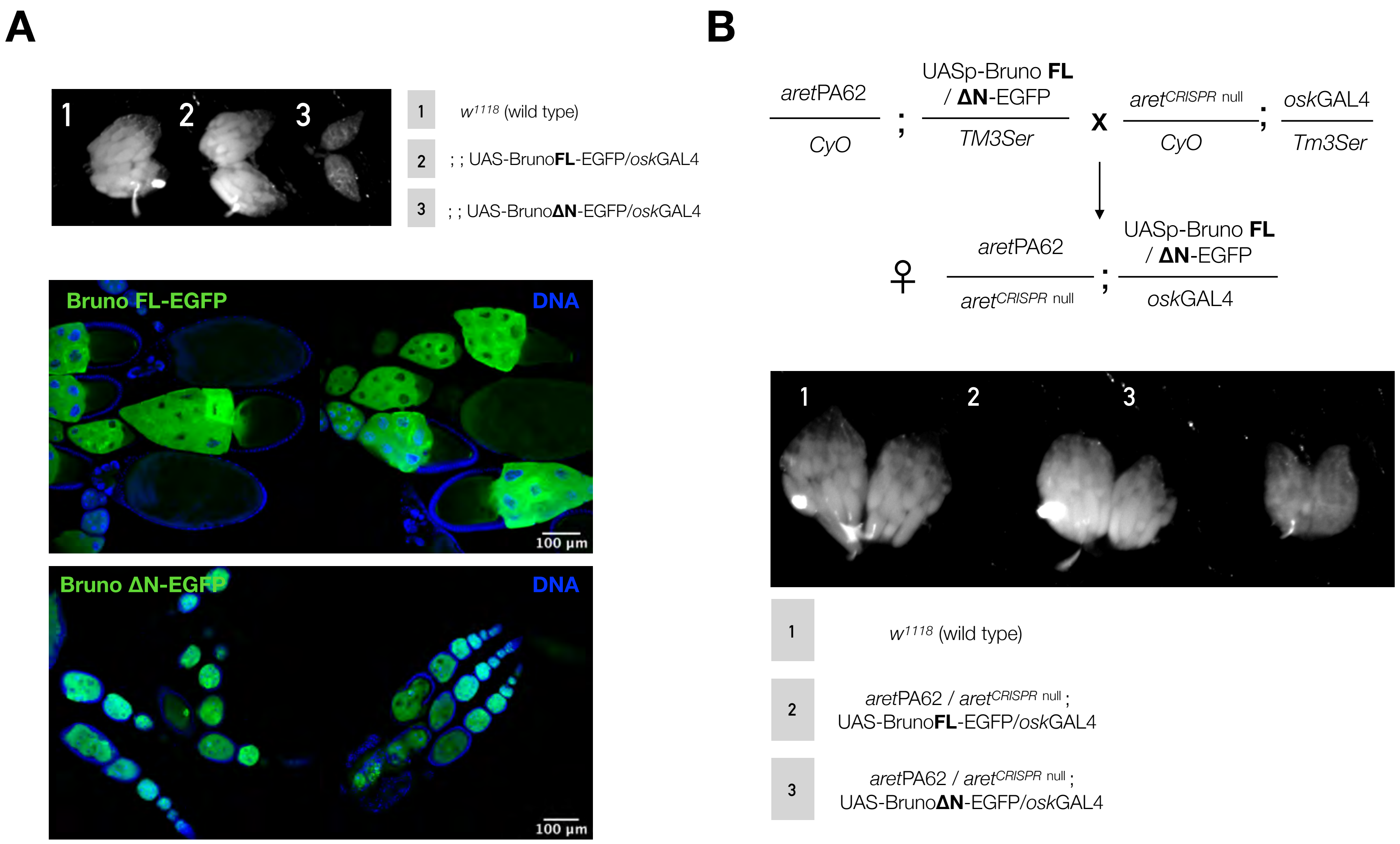

**Figure S7, Related to Figure 4**

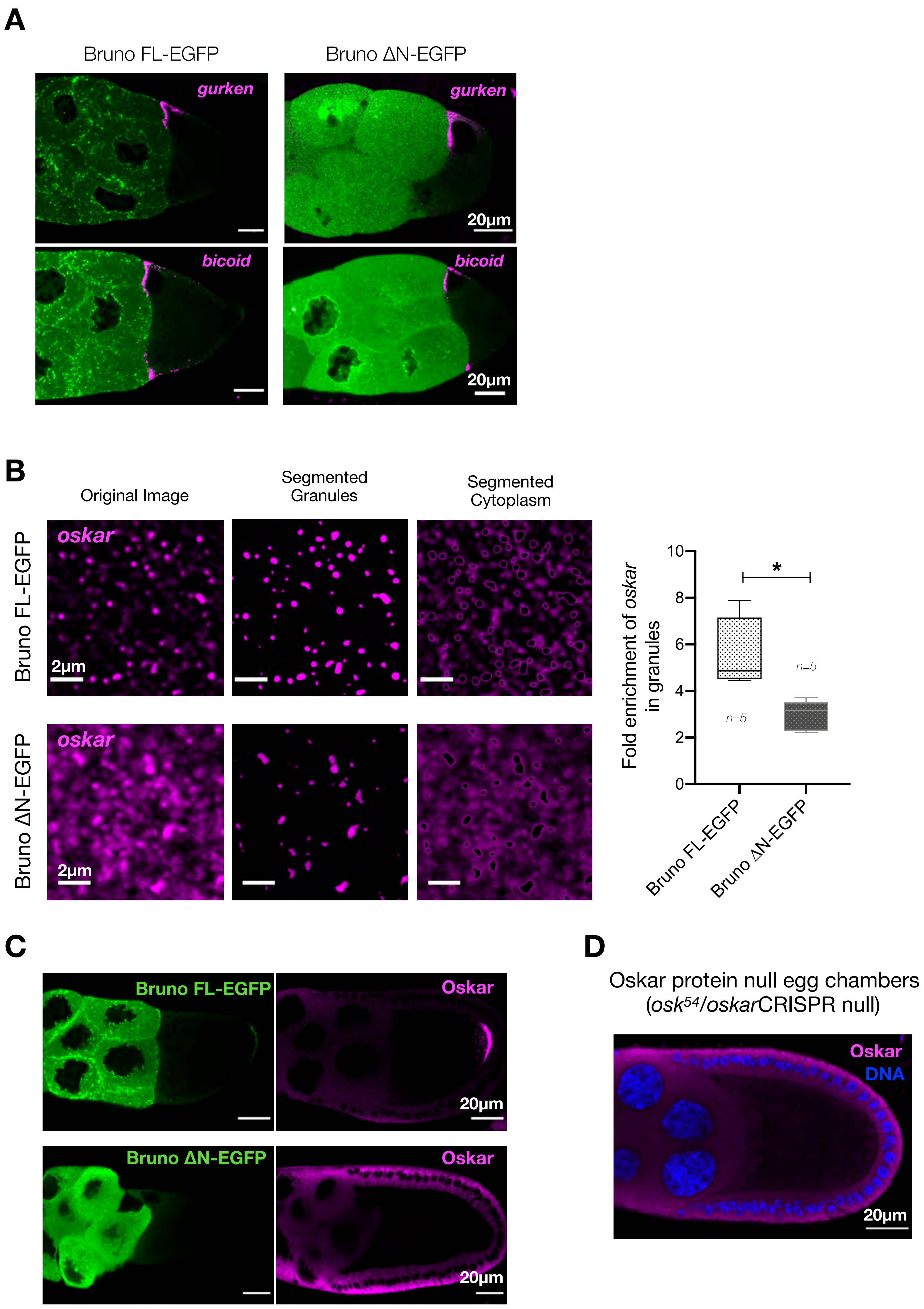

**Figure S8, Related to Figure 4**

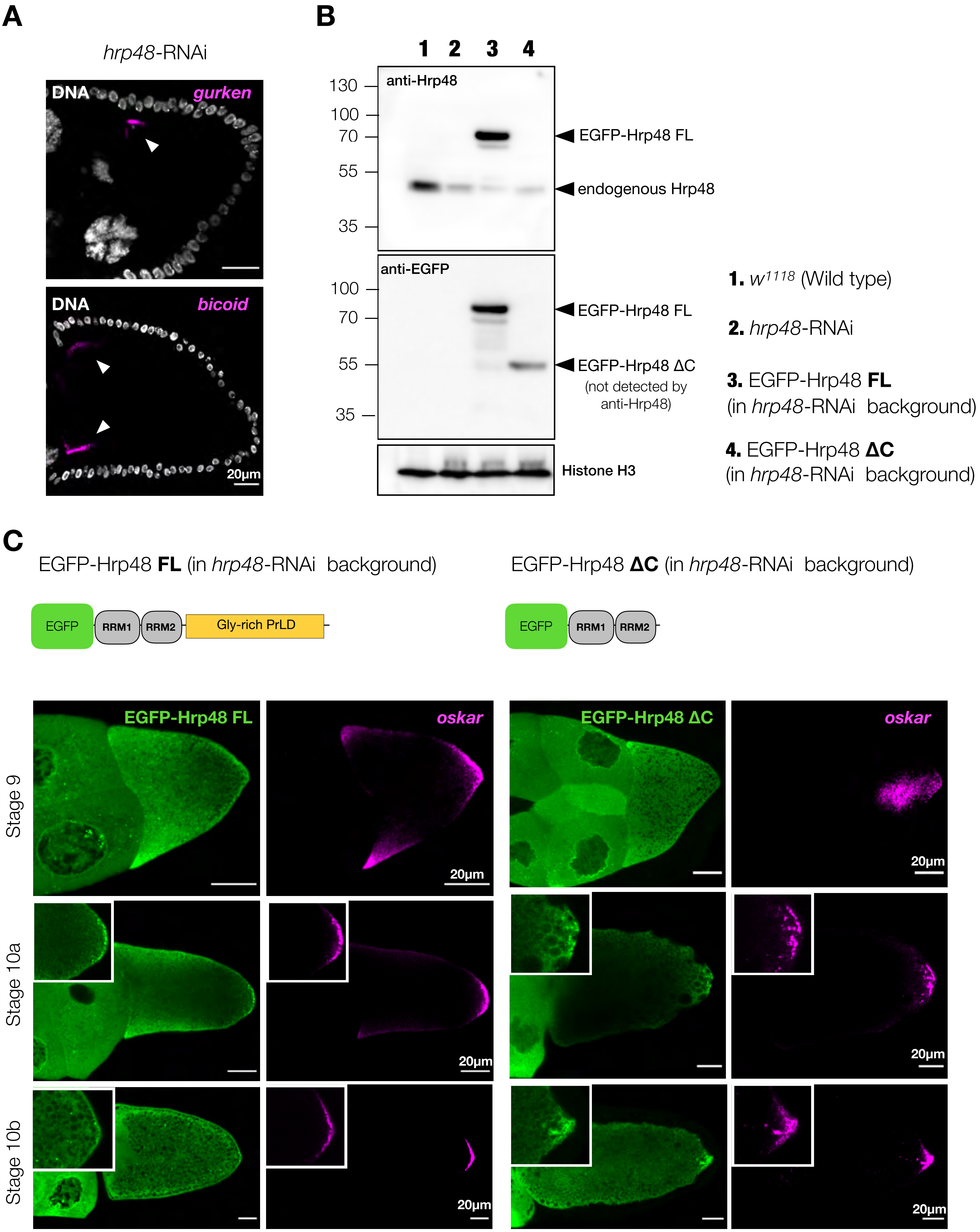

**Figure S9, Related to Figure 5**

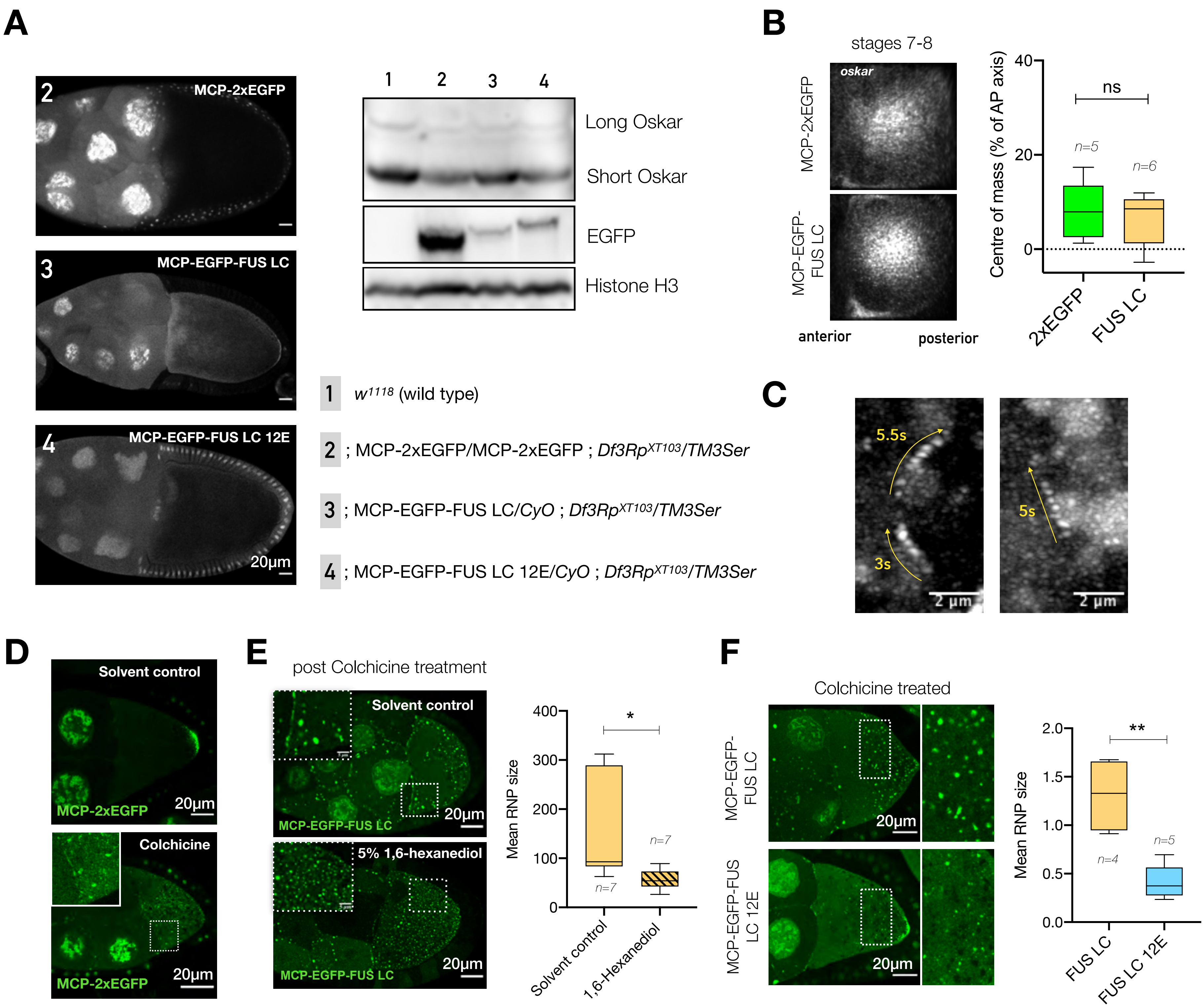

**Figure S10, Related to Figure 6**

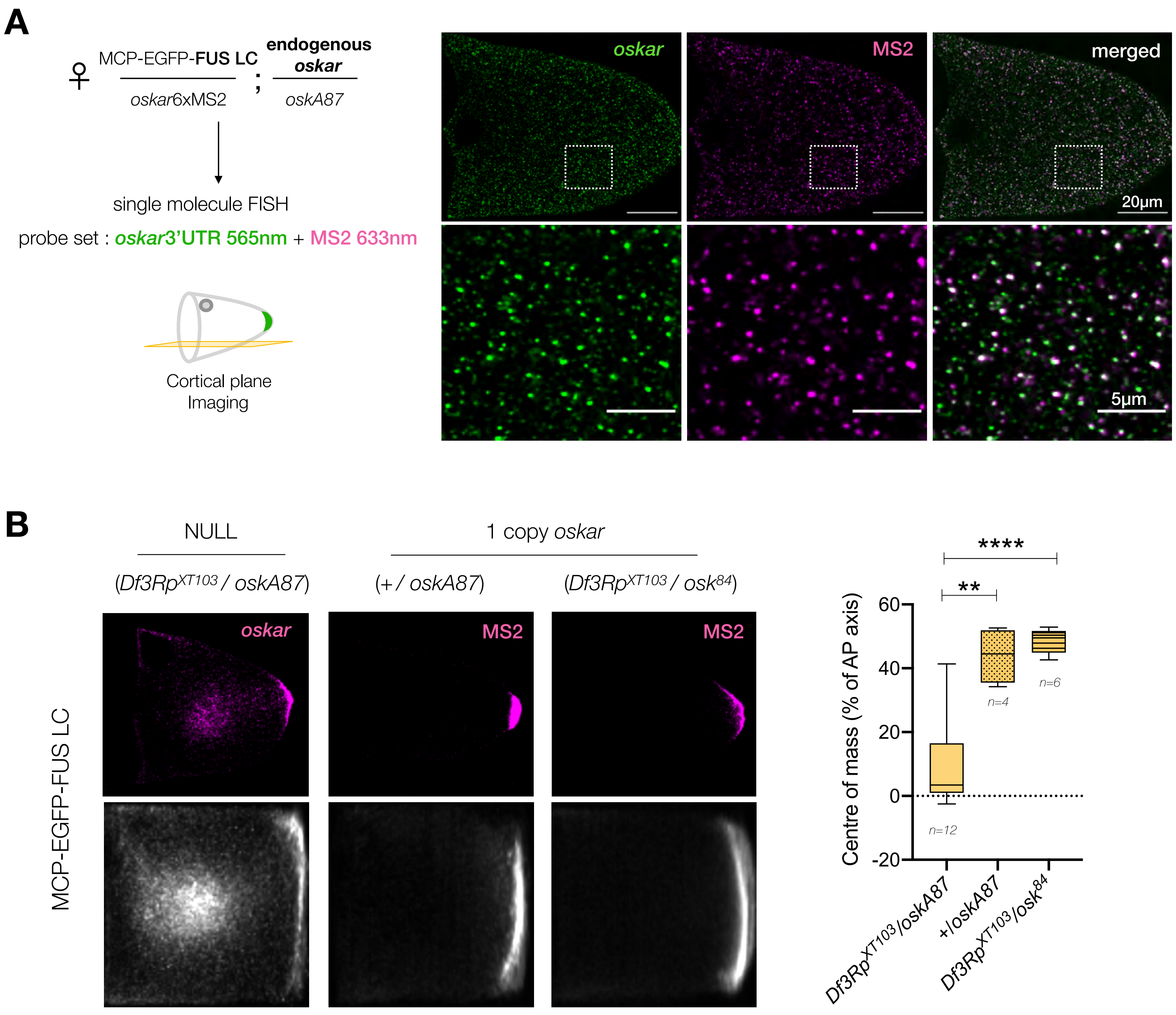

**Figure S11, Related to Figure 6**

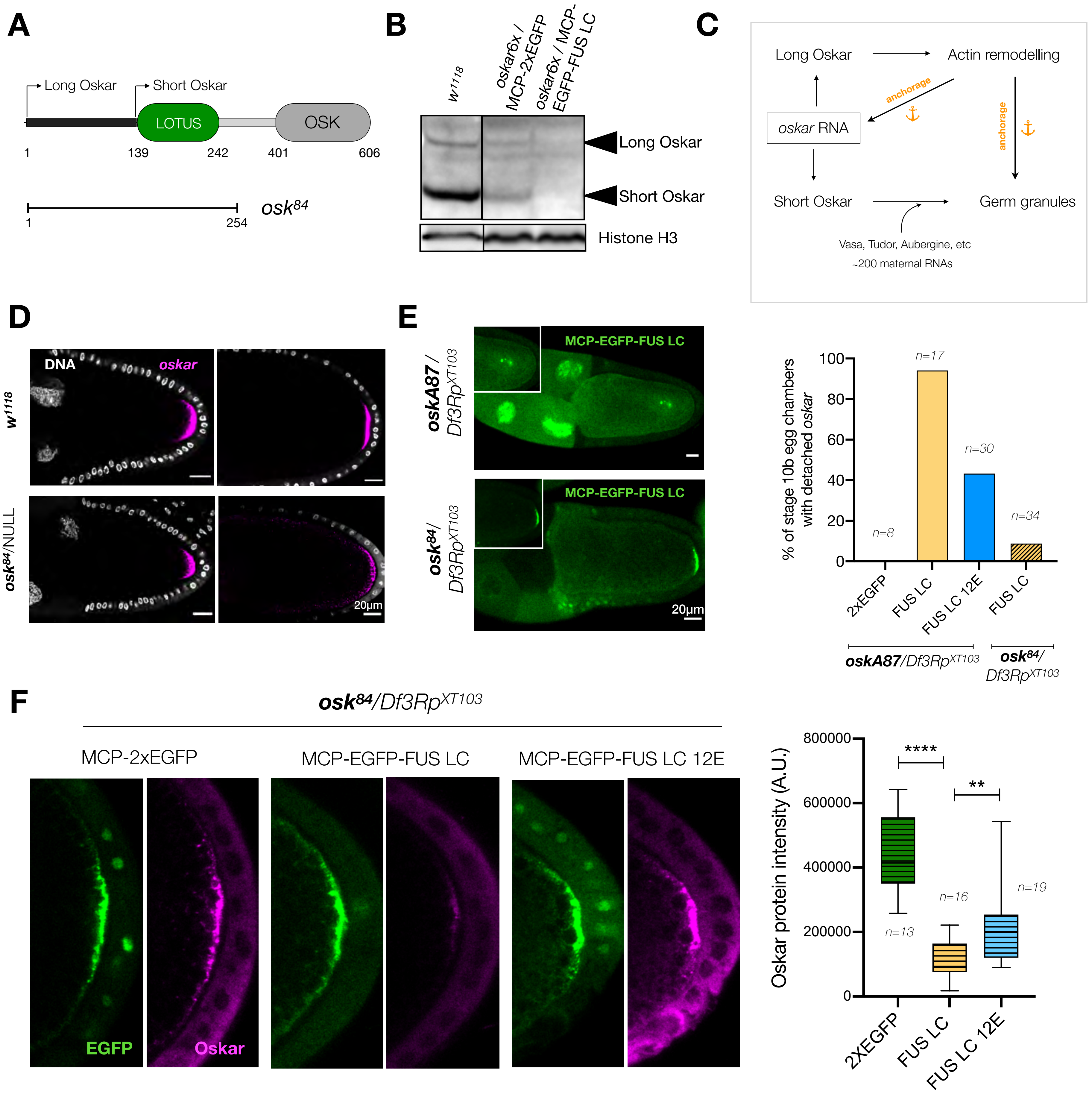

**Figure S12, Related to Figure 7**

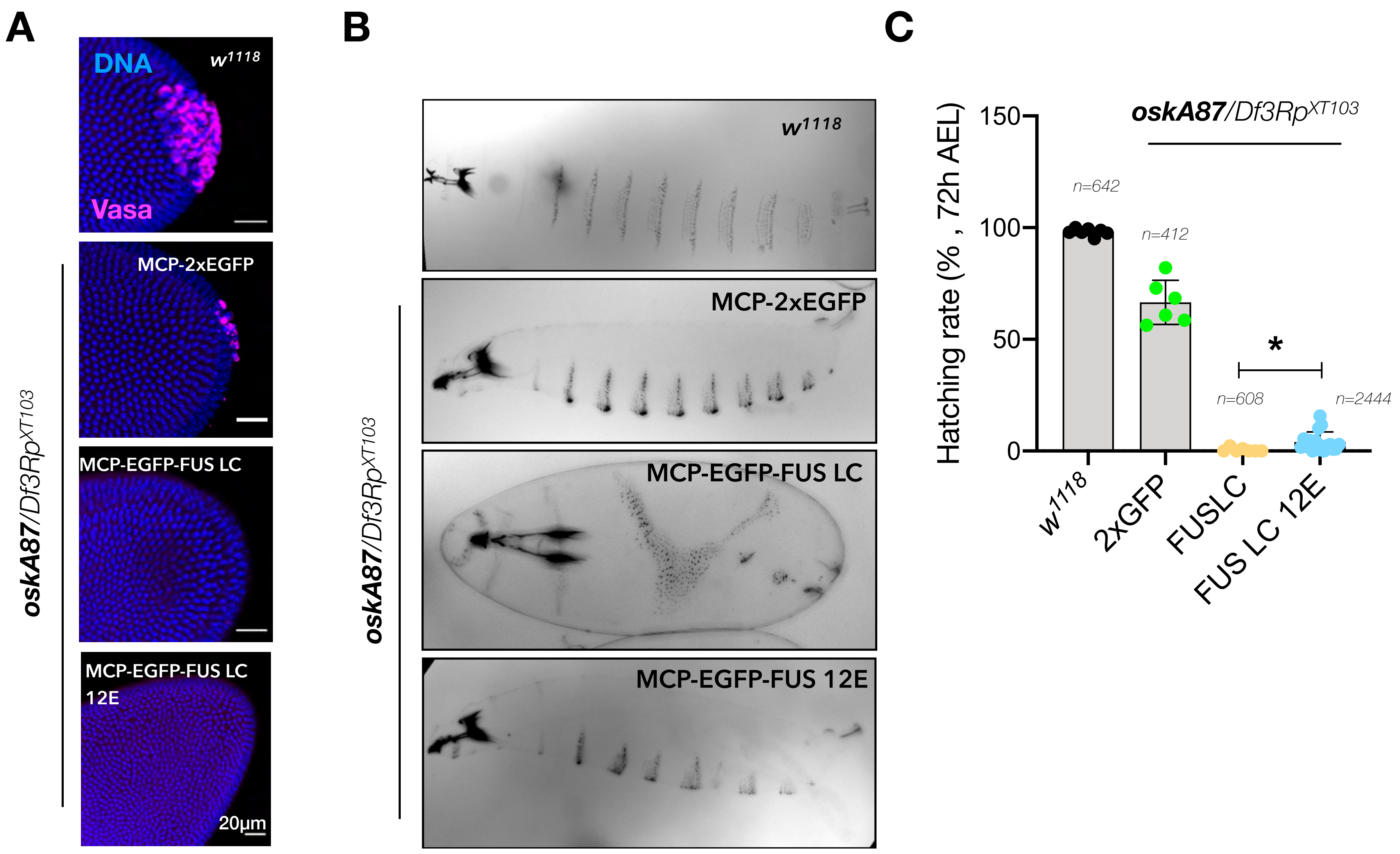

**Figure S13, Related to Figure 7**

**Supplemental Information titles and Legends**

**Figure S1, Related to Figure 2**

(A) *oskar* mRNA association with 3 *bona fide* granule proteins. smFISH for *oskar* mRNA (magenta) and the respective proteins (green). A maximum intensity projection of a Z-stack of 1 µm is shown for nurse cell and oocyte (cortical plane). For Bruno, BrunoFL-EGFP was conditionally expressed in the germline in an endogenous Bruno-deficient genetic background (*aret*PA62/*aret*$^{CRISPR\text{null}}$). For Hrp48, EGFP-Hrp48FL was expressed in the germline in an Hrp48 RNAi background (refer to Figure S9B for knockdown efficiency). For PTB, a homozygous EGFP-trap line was used in which both alleles of PTB bear the GFP insertion. Insets are marked with a dotted white line and are also shown in Figure 2A.

(B) Domain architecture of the 3 core *oskar* granule proteins with disorder prediction in IUPred (Meszaros et al., 2018).

**Figure S2, Related to Figure 2**

(A) Coomassie stained SDS-PAGE gel with the 3 granule proteins purified from Sf-21 insect cells.

(B) EMSA demonstrating the intrinsic affinities of the purified core granule proteins for *oskar* 3'UTR RNA. 50 nM atto633-labelled *oskar* 3'UTR was used with the indicated concentrations of the proteins and the reaction was resolved on an agarose gel.

(C-D) Schematic representation of the *in vitro* condensate assembly assay for Bruno and Hrp48. 10 µM of tagged protein incubated with 1 U of SumoStar protease for 30 min at room temperature followed by SDS-PAGE or imaging. SDS-PAGE shows the efficiency of tag cleavage in assay buffer with 300mM NaCl; * indicates the cleaved protein band (C). SumoStar tag cleavage does not induce condensate formation in 300 mM NaCl buffer. Exchange to 150 mM NaCl buffer triggers phase separation of Bruno and Hrp48 (D).

(E) 100 nM *oskar* 3'UTR-atto633 (red) does not self-assemble into condensates in absence of protein under the same conditions.

(F) Molecular crowding promotes condensate formation of *oskar* 3'UTR with PTB. 10 µM RFP-PTB (green) is soluble in 150 mM NaCl buffer in absence or presence of *oskar* 3'UTR (magenta). Addition of 5% (w/v) PEG-4000 induces formation of spherical condensates of PTB alone and with *oskar* 3'UTR.

(G) Condensates formed with 10 µM Bruno-EGFP or Hrp48-EGFP (green) and 100 nM *oskar* 3'UTR do not fuse and relax like liquid droplets. Also refer to movies S4-S6.

(H) Condensates assembled with 100 nM *oskar* 3'UTR (unlabeled) and 10 µM of Bruno or Hrp48 (green); 8µM hFUS-EGFP (without RNA; green) with 10% PEG-4000 was used as a control for a well characterized FRAP behavior in liquid condensates. FRAP movie snapshots at indicated time points with bleached region of interest (ROI) marked with a dotted circle. Quantification is provided in Figure 2D. For time lapse videos refer to Movie S7.

**Figure S3, Related to Figure 2**

(A) Cryo-EM image of Bruno-*oskar* 3'UTR condensates deposited on a holey carbon EM grid. Leftmost panel shows a grid map with varying ice thickness and yellow dotted box represents grid squares enlarged on the right. Within an individual square, deposited condensates are indicated by black arrows. Small condensates deposited in holes of the holey carbon film and therefore amenable to tilt series acquisition are marked with white arrows, whereas black arrow marks a condensate on the support film that is too thick to be imaged.

(B) Cryo-EM images of enlarged view of holes within a grid square with Bruno-*oskar* 3'UTR (left) and Hrp48-*oskar* 3'UTR (right) condensates. Left panel shows a cluster of spherical condensates too thick for acquiring tilt-series; inset shows an enlarged view of spherical condensates. Right panel shows two spherical isolated condensates; inset shows an enlarged view of one that is optimal for tilt-series acquisition.

**Figure S4, Related to Figure 3**

(A) Quantification of *in vivo* protein concentrations per granule. For protein quantifications, GFP-trap lines of Bruno and PTB were used and absolute concentrations calculated based on a calibration curve of recombinant EGFP imaged under identical conditions in the same imaging session. Numbers in the histogram refer to mean number of granules grouped under the indicated range of concentration.

(B) For *in vivo oskar* RNA concentrations per granule, $w^{1118}$ (wild type) egg chambers were stained for *oskar* by smFISH and *oskar* copy number per granule in the oocyte compartment was calculated. Granule volume obtained from 3D STED experiments was plotted and absolute molar concentration of *oskar* RNA per granule was then derived based on average granule volume. Numbers in the histogram refers to number of granules grouped under the indicated range of volume.

(C) Representative light microscopy single plane confocal images of experimental conditions used for cryo-electron tomography in Figure 3B. Images have been acquired and processed independently.

(D) Condensates with Bruno alone preclude incorporation of *oskar* 3'UTR. Condensates were assembled with Bruno alone in 150 mM NaCl assay buffer and 10 nM atto633 labelled *oskar* 3'UTR RNA was added at 30 min of condensate ageing. In case of 30 min time point condensates exclude RNA. Note that new condensates formed after RNA addition show colocalization of the RNA and protein (marked by *).

(E) Plot of mRNA intensity versus granule volume of *oskar* RNP granules measured by 3D STED on $w^{1118}$ egg chambers probed for *oskar* mRNA by smFISH. Intensity of *oskar* mRNA signal (top plot) was normalized by granule volume (B) to derive RNA concentration per granule, which does not increase with increase in granule volume.

**Figure S5, Related to Figure 4**

(A) Western blot depicting knockdown of PTB upon RNAi driven by *oskar*GAL4 driver in the germline.

(B-C) Posterior localization of *oskar* as detected by smFISH (B) as well as translation of Oskar protein (C) is unaffected upon PTB knockdown.

**Figure S6, Related to Figure 4**

(A) Sequence alignment of amino acids 1-179 of *Drosophila melanogaster* Bruno and orthologs in other Drosophilids.

(B) Expression of Bruno FL-EGFP and ∆N-EGFP in Schneider cells (S2R+). Note that Schneider cells do not express *oskar* mRNA.

(C) *in vitro* reconstitution of 10 µM Bruno FL-EGFP and ∆N-EGFP in 150 mM NaCl containing assay buffer.

**Figure S7, Related to Figure 4**

(A) Overexpression of EGFP-tagged FL and ∆N Bruno in the germline driven by *oskar*GAL4. Ovary morphology indicates atrophic ovaries upon overexpression of ∆N-EGFP. Protein is in green and nuclei are stained with DAPI.

(B) Scheme of the genetic cross carried out for conditional expression of EGFP-tagged Bruno FL and ∆N in the female germline in a Bruno-deficient background (*aret*PA62/$aret^{CRISPR\ null}$). Morphology of ovaries of the indicated genotype is shown along with wild type ($w^{1118}$).

**Figure S8, Related to Figure 4**

(A) Localization of *bicoid* and *gurken* is not affected upon expression of EGFP-tagged Bruno FL and ∆N in the female germline in a Bruno-deficient background (*aret*PA62/$aret^{CRISPR\ null}$). smFISH detected *gurken* (magenta) localizing correctly at the dorso-anterior corner and *bicoid* (magenta) at the anterior margin of the oocyte in mid oogenesis.

(B) Granular morphology of *oskar* RNPs is lost significantly in ∆N-EGFP. Representative single confocal plane of ooplasm with *oskar* (magenta) labelled by smFISH. Segmentation of granules and ooplasm depicted. Quantification of fold enrichment of RNA inside granules (ratio of mean intensity inside granules to mean total intensity). Error bars represent SD. Unpaired Student's t-tests were used for comparisons. Significance level: * < 0.05.

(C-D) Translation of Oskar protein is not detected upon expression of ∆N. Immunostaining for Oskar protein (magenta) in early stage 10 egg chambers detected Oskar protein in Bruno FL, but not ∆N expressing egg chambers. In case of ∆N, signal (magenta) from the periphery of egg chamber is background fluorescence (C) and is also detected in Oskar protein null flies (D).

**Figure S9, Related to Figure 5**

(A) Localization of maternal RNAs *gurken* and *bicoid* (magenta) is not affected upon *hrp48*-RNAi. Maximum intensity projection of a Z-volume of 5 µm.

(B) Western blot of ovaries from flies of the indicated genotypes. The blot probed with anti-Hrp48 antibody has been stripped and re-probed with anti-GFP, as anti-Hrp48 failed to detect the truncated ∆C version. Histone H3 serves as loading control.

(C) Constructs of Hrp48 FL and ∆C used for fly transgenesis. The images are from flies expressing the EGFP-tagged proteins in the *hrp48*-RNAi background. Representative confocal images of egg chambers of stages 9, 10a and 10b shown with Hrp48 variants (green) and *oskar* (magenta) detected by smFISH. Insets show enlarged version of the posterior pole.

**Figure S10, Related to Figure 6**

(A) Representative images of stage 10 egg chambers with EGFP signal in grayscale; respective genotypes are indicated. Western blot of ovaries from the indicated genotypes shows transgene expression levels (anti-EGFP antibody) and Oskar protein. Histone H3 serves as loading control.

(B) Mean *oskar* RNA signal (grayscale) from smFISH data from several stage 7-8 oocytes; anterior to the left. Position of the *oskar* center of mass relative to the geometric center of the oocyte (dotted horizontal line) along the AP axis is indicated. Error bars represent SD. Unpaired Student's t-tests were used for comparisons. ns: non-significant.

(C) Maximum Z-projections of selected regions of Movie S15 (MCP-EGFP-FUS LC in grayscale) showing fast, directed tracks marked with yellow arrows with all three depicted particles having a velocity > 0.5 µm/sec.

(D) Colchicine treatment on isolated ovaries in 2xEGFP tethered condition.

(E) Treatment of egg chambers with 5% 1,6-Hexanediol for 15 min after 2 h of colchicine treatment dissolves the large spherical assemblies partially. Quantification of granule size shows a significant reduction in 1,6-HD treated samples. Error bars represent SD. Unpaired Student's t-tests were used for comparisons. Significance level: * < 0.05.

(F) Colchicine treatment of stage 8-9 egg chambers induced the formation of large granules in case of FUS LC, which is significantly reduced in 12E. Error bars represent SD. Unpaired Student's t-tests were used for comparisons. Significance level: ** < 0.01.

**Figure S11, Related to Figure 6**

(A) *oskar*6XMS2 mRNA and endogenous *oskar* transcripts co-package into the same granules. Two-color smFISH of egg chambers expressing one endogenous genomic *oskar* and the *oskar* 6xMS2 transgene with atto-565 probes against *oskar* (green) and atto-633 probes against MS2 loops (magenta). Bottom panels are enlarged from boxed regions.

(B) Dilution of FUS LC per granule by an endogenous copy of *oskar* mRNA recuses the transport defects. *oskar* distribution in representative stage 9 egg chambers detected by smFISH (magenta). Position of the *oskar* center of mass relative to the geometric center of the oocyte (dotted horizontal line) along the AP axis in indicated genetic backgrounds. *n* denotes the number of oocytes analysed. Error bars represent SD. Unpaired Student's t-tests were used for comparisons. Significance levels: **<0.01, ****<0.0001.

**Figure S12, Related to Figure 7**

(A) Schematic representation of Oskar protein domain architecture indicating the start sites of the Long and Short isoforms. Nonsense mutant $osk^{84}$ putatively expresses 254 residues from the N-terminus.

(B) Western blot for Oskar protein confirms loss of translation upon FUS LC tethering. Arrows mark the two isoforms of Oskar protein. Note that the reduction in Oskar protein levels in case of MCP-2xEGFP compared to wild type is due to the *oskar* RNA null (*oskA87*/$Df3Rp^{XT103}$) background of the flies.

(C) Schematic representation of multiple interdependent functions of Oskar protein isoforms in actin remodeling, anchoring and organization of the germ plasm (Tanaka and Nakamura, 2011).

(D) Anchoring of *oskar* RNPs is rescued in females expressing an $osk^{84}$ allele; NULL indicates the other chromosome: *oskar* CRISPR RNA null allele. smFISH for *oskar* mRNA (magenta) on egg chambers of the indicated genotypes shows *oskar* anchoring in late stage 9 (left) and stage11-12 (right) egg chambers.

(E) Anchoring defects are appreciably rescued in $osk^{84}$/$Df3Rp^{XT103}$ background. Representative images of stage 10b egg chambers expressing MCP-EGFP-FUS LC (green) in the indicated genetic backgrounds. Quantification of *oskar* detachment phenotype from images of stage 10b egg chambers expressing the indicated transgenes in absence or presence of anchoring provided in trans. *n* denotes the number of egg chambers analysed.

(F) Immunostaining of egg chambers for Oskar protein (magenta) upon provision of anchoring *in trans* by the $osk^{84}$ allele ($osk^{84}/Df3Rp^{XT103}$). The EGFP signal in green confirms the rescue of anchoring. All images shown (and used for quantification) were acquired using identical microscope settings and representations are contrast-matched. Quantification of the Oskar signal intensity from the posterior of several egg chambers confirms the reduction of translation upon FUS LC tethering and partial translation using the FUS 12E construct. Note that FUS LC panel is also shown in Figure 7C. Error bars represent SD and *n* denotes number of analysed oocytes. Unpaired Student's t-tests were used for comparisons. Significance levels: ** < 0.01, **** < 0.0001.

**Figure S13, Related to Figure 7**

(A) Formation of the germline is impaired upon FUS LC and 12E tethering in an *oskar* RNA null background ($oskA87/Df3Rp^{XT103}$). Reduction of pole cell numbers is noted in 2xEGFP tethering compared to wild type ($w^{1118}$). Pole cells at the posterior of embryos at NC14 are identified by Vasa (magenta) immunostaining. Nuclei stained with DAPI (blue).

(B) Cuticles of embryos reveal severe patterning defects upon FUS LC tethering. Anterior is to the left and ventral faces the bottom.

(C) Quantification of the hatching rates of eggs from females expressing the indicated transgene in an *oskar* RNA null background ($oskA87/Df3Rp^{XT103}$). Number of eggs scored per genotype is depicted in the graph. Note that data for $w^{1118}$ are also shown in Figure 7F. Error bars represent SD and *n* denotes number of analysed eggs. Unpaired Student's t-tests were used for comparisons. Significance level: * < 0.05.

**Supplementary Movies**

**Movie S1.** 3D STED volume of *oskar* RNP granules (yellow) in the oocyte center, Related to Figure 1

**Movie S2.** Fluorescence time-lapse movie of *oskar* granules labelled with MCP-EGFP (grayscale) close to the posterior pole, Scale 2 μm, Related to Figure 1

**Movie S3.** Fluorescence time-lapse movie of *oskar* granules labelled with MCP-EGFP (grayscale) at the posterior cortex, Scale 2 μm, Related to Figure 1

**Movie S4.** Time-lapse movie of condensates with 10 μM Bruno-EGFP (green) and 100 nM *oskar* 3'UTR, showing the non-dynamic behavior of the condensates. Related to Figure 2

**Movie S5.** Time-lapse movie of condensates with 10 μM Hrp48-EGFP (green) and 100 nM *oskar* 3'UTR showing the non-dynamic behavior of the condensates. Related to Figure 2

**Movie S6.** Time-lapse movie of condensates with 8 μM hFUS-EGFP (green) and 10% PEG-4000, showing the dynamic fusion behavior of the condensates. Related to Figure 2

**Movie S7.** FRAP time lapse movies of condensates formed with 10 μM Bruno-EGFP or Hrp48-EGFP (green) and 100 nM *oskar* 3'UTR. FUS condensates are formed with 8 μM FUS-EGFP (green) and 10% PEG-4000, Scale bar 5μm, Related to Figure 2

**Movie S8.** Tomographic reconstruction of condensate formed with 10 μM Bruno-EGFP and 100 nM *oskar* 3'UTR, Scale bar 100 nm, each slice is 4 nm thick, volume was gaussian filtered with kernel 3. Related to Figure 2

**Movie S9.** Tomographic reconstruction of condensate formed with 10 μM Bruno-EGFP and 100 nM full length *oskar* mRNA, Scale bar 100 nm, each slice is 4nm thick, volume was gaussian filtered with kernel 3. Related to Figure 2

**Movie S10.** Tomographic reconstruction of condensate formed with 10 µM Hrp48-EGFP and 100 nM *oskar* 3'UTR, Scale bar 100 nm, each slice is 4 nm thick, volume was gaussian filtered with kernel 3. Related to Figure 2

**Movie S11.** Tomographic reconstruction of Bruno-*oskar* 3'UTR condensate (with 400 nM labelled *oskar* 3'UTR added at 0 min), Scale bar 100 nm, each slice is 4 nm thick, volume was gaussian filtered with kernel 3. Related to Figure 3

**Movie S12.** Tomographic reconstruction of Bruno-*oskar* 3'UTR condensate (with 400 nM labelled *oskar* 3'UTR added at 30 min), Scale bar 100 nm, each slice is 4 nm thick, volume was gaussian filtered with kernel 3. Related to Figure 3

**Movie S13.** Tomographic reconstruction of Bruno-*oskar* 3'UTR condensate (2 µM RFP-PTB added at 30 min), Scale bar 100 nm, each slice is 4 nm thick, volume was gaussian filtered with kernel 3. Related to Figure 3

**Movie S14.** A 3.2 µm Z-stack of nurse cells showing Bruno FL-EGFP (green) and *oskar* mRNA FISH (magenta) association on track-like structures, Related to Figure 4

**Movie S15.** Directed transport of FUS-tethered *oskar* granules (grayscale) near posterior pole, Related to Figure 6

**Movie S16.** Fusion of large FUS-tethered *oskar* granules (green) in oocyte upon colchicine treatment. A cropped region is presented in Figure 6E, Related to Figure 6

**Movie S17.** Fusion of FUS-tethered *oskar* granules (grayscale) into larger structures at the posterior pole. Inset shows an enlarged view, Related to Figure 7

**STAR Methods**

**Key Resources table**

| REAGENT/RESOURCE | SOURCE | IDENTIFIER |
|---|---|---|
| **Antibodies** | | |
| Rabbit anti-Oskar (1:3000; WB & IF) | In house | -- |
| Rabbit anti-Hrp48 (1:2000; WB) | In house | -- |
| Rabbit anti-PTB (1:2000; WB) | In house | -- |
| Rabbit anti-EGFP (1:5000; WB) | Torrey Pines Biolabs | Cat# TP401 |
| Rabbit anti-Histone H3 (1:2500; WB) | Abcam | Cat# ab1791 |
| Rat anti-Vasa (1:500; IF) | In house | -- |
| **Chemicals** | | |
| 1,6-Hexanediol | Sigma | Cat# 240117 |
| Colchicine | Sigma | Cat# C9754 |
| Insulin | Sigma | Cat# I9278 |
| Fetal Bovine Serum | Life Technologies | Cat# 10082147 |
| Schneider's Drosophila medium | Life Technologies | Cat# 21720024 |
| SumoStar protease | Life Sensors | Cat# 4110 |
| PEG-4000 | Thermo Scientific | Cat# EL0011 |
| ATP | Thermo Scientific | Cat# R0441 |
| RNase A | Thermo Scientific | Cat# EN0531 |
| Tetramethyl Rhodamine BSA | Thermo Scientific | Cat# A23016 |
| Shandon ImmuMount | Thermo Scientific | Cat# 9990402 |
| **Critical Commercial assays** | | |
| Effectene Transfection reagent | Qiagen | Cat# 301425 |
| MEGAscript T7 transcription kit | Thermo Scientific | Cat# AMB13345 |

| | | |
|---|---|---|
| StrepTrap HP | Merck | Cat# GE28-9075-47 |
| HisTrap HP | Merck | Cat# GE17-5248-01 |
| HiLoad 16/600 Superdex 200pg | Merck | Cat# GE28-9893-35 |
| **Plasmids** | | |
| pFastBAC 1-mRFP-PTB | This study | -- |
| pCoofy63-BrunoFL-EGFP | This study | -- |
| pCoofy63-BrunoΔN -EGFP | This study | -- |
| pCoofy63-Hrp48FL-EGFP | This study | -- |
| pCoofy63-Hrp48ΔC -EGFP | This study | -- |
| UASp-attB-K10-Bruno FL-EGFP | This study | -- |
| UASp-attB-K10-Bruno ΔN-EGFP | This study | -- |
| pU6-BbsI-chiRNA | https://flycrispr.org | Addgene Cat# 45946 |
| pHD-scarless dsRED | https://flycrispr.org | Addgene Cat# 64703 |
| UASp-attB-K10-EGFP-Hrp48FL | This study | -- |
| UASp-attB-K10-EGFP-Hrp48ΔC | This study | -- |
| pHsp83-NLS-HA-tdMCP-2xEGFP | This study | -- |
| Dendra2-FUS WT | (Patel et al., 2015) | Gift from Hyman lab |
| pHsp83-NLS-HA-tdMCP-EGFP-FUS LC | This study | -- |
| MBP-FUS FL-12E | (Monahan et al., 2017) | Addgene Cat# 98652 |
| pHsp83-NLS-HA-tdMCP-EGFP-FUS LC 12E | This study | -- |
| pActin5C-Bruno FL-EGFP | This study | -- |
| pActin5C-Bruno ΔN -EGFP | This study | -- |

| | | |
|---|---|---|
| **Oligonucleotide probes** | | |
| FISH probes are listed in Table S1 | (Gaspar et al., 2017b) | -- |
| **Fly strains** | | |
| D. melanogaster: *w*$^{1118}$ | BDSC | 3605 |
| D. melanogaster: *oskar6XMS2*/*CyO* | Gift from Elizabeth Gavis lab (Lin et al., 2008) | -- |
| D. melanogaster: *oskar6XMS2:MCP-EGFP/Tm3Sb* | Gift from Elizabeth Gavis lab (Lin et al., 2008) | -- |
| D. melanogaster: *mRFP-Nup107* | Gift from Martin Beck lab (Hampoelz et al., 2019) | -- |
| D. melanogaster: | BDSC | 51324 |
| D. melanogaster: y[1] M{vas-int.Dm}ZH-2A w[*]; PBac{y[+]-attP-3B}VK00033 | BDSC | 24871 |
| D. melanogaster: vas-phi-ZH2A, PBac{y[+]-attP-9A}VK00018 | BDSC | 9736 |
| D. melanogaster: *aret*CRISPR-dsRED/*CyO* | This study | -- |
| D. melanogaster: *If*/*CyO*; UASp-BrunoFL-EGFP/*Tm3ser* | This study | -- |
| D. melanogaster: *If*/*CyO*; UASp-Bruno∆N-EGFP/*Tm3ser* | This study | -- |
| *bruno*-RNAi; D. melanogaster: *P{TRiP.HMC02374}attP2* | BDSC | 44483 |

| | | |
|---|---|---|
| *hrp48*-RNAi; D. melanogaster: *P{TRiP.JF01478}attP2* | BDSC | 31685 |
| *ptb*-RNAi; D. melanogaster: *P{TRiP.GLV21034}attP2* | BDSC | 35669 |
| D. melanogaster: *If*/*CyO*; UASp- EGFP-Hrp48FL- /*Tm3ser* | This study | -- |
| D. melanogaster: *If*/*CyO*; UASp- EGFP-Hrp48ΔC- /*Tm3ser* | This study | -- |
| *PTB-EGFP trap;* D. melanogaster: *w-;Bl/CyO;GFP-PTB/TM6b* | Ephrussi lab (Besse et al., 2009) | - |
| *Bruno-EGFP trap;* D. melanogaster: *w-; GFP-Bruno/CyO* | BDSC | 60144 |
| D. melanogaster: pHsp83-MCP-2xEGFP/*CyO* | This study | -- |
| D. melanogaster: pHsp83-MCP-EGFP-FUS LC/*CyO* | This study | -- |
| D. melanogaster: *If*/*CyO*; pHsp83-MCP-EGFP-FUS LC 12E/*Tm3ser* | This study | -- |
| D. melanogaster: *oskar*GAL4/*Tm3sb* | BDSC | 44242 |
| D. melanogaster: *oskar A87* | | |
| D. melanogaster: $oskar^{attP,3P3GFP}$ | (Gaspar et al., 2017a) | |
| D. melanogaster: $oskar^{84}$ | | |
| D. melanogaster: $Df3Rp^{XT103}$ | | |
| **Software** | | |
| Fiji | - | https://fiji.sc |

| | | |
|---|---|---|
| xsPT Fiji plugin | (Gaspar and Ephrussi, 2017) | https://github.com/Xaft/xs/blob/master/_xs.jar |
| Cort Analysis Fiji plugin | (Gaspar et al., 2014) | - |
| Imaris | - | https://imaris.oxinst.com |
| Huygens Essential | - | https://svi.nl/Huygens-Deconvolution |
| FRAP Analyser | - | https://github.com/ssgpers/FRAPAnalalyser |
| PLAAC | (Lancaster et al., 2014) | http://plaac.wi.mit.edu/ |
| IUPred | (Meszaros et al., 2018) | https://iupred2a.elte.hu |
| IMOD v.4.9 | - | https://bio3d.colorado.edu/imod/ |
| TOM package | (Nickell et al., 2005) | MATLAB (MathWorks) |
| SerialEM v3.7.2 | (Mastronarde, 2005) | - |
| WARP | (Tegunov and Cramer, 2019) | http://www.warpem.com/warp/ |
| Prism 8.0 | GraphPad | https://www.graphpad.com/ |
| JalView 2..11.1.3 | - | http://www.jalview.org |

*WB: Western Blotting; IF: Immunofluorescence; td:tandem.

**Lead Contact and Materials Availability**

Further information and requests for resources and reagents should be directed to and will be fulfilled by the Lead contact Anne Ephrussi. Information about and request for tomography data should be directed to and will be fulfilled by Julia Mahamid. All unique materials and reagents generated in this study are available from the lead contact with a completed material transfer agreement.

**Experimental Model and Subject Details**

*Drosophila melanogaster* fly stocks were maintained at 25°C on standard cornmeal agar. 3-6 day old female flies with typically half as many male flies were transferred to vials with fresh yeast 24h before experiments.

#### Method Details

##### Fly stocks

The following fly strains were used in the study: *w*$^{1118}$ ; *oskar6xMS2/CyO (gift from Elizabeth Gavis); oskar6xMS2:MCP-EGFP/Tm3Sb (gift from Elizabeth Gavis); mRFP-Nup107 (gift from Martin Beck); hrp48-RNAi: P{TRiP.JF01478}attP2; bruno-RNAi: P{TRiP.HMC02374}attP2; ptb-RNAi: P{TRiP.GLV21034}attP2 ; PTB-EGFP trap: w-;Bl/CyO;GFP-PTB/TM6b.*

##### Generation of transgenic flies

*P-element mediated germline transformation*

Transgenic flies expressing MCP-2xEGFP/MCP-EGFP-FUS LC/MCP-EGFP-FUS LC 12E were generated by P-element transformation. Briefly, 2xEGFP, FUS LC (gift from Tony Hyman) and FUS LC 12E (Addgene) were cloned under the Hsp83 promoter and P-element transgenesis performed in *w*$^{1118}$ flies using standard procedures (Rubin and Spradling, 1982). After eye color based screening for positives using the truncated *mini-white* marker gene, transgenes were balanced on the respective chromosomes (*CyO* or *Tm3Ser*) to establish stable stocks. For FUS LC 12E, inserted on chromosome III, recombination with an *oskar* CRISPR null line expressing EGFP under the 3xP3 promoter (*oskar*$^{attP,3P3GFP}$ ; Gasper et al., 2017) was carried out.

*Bruno CRISPR knock out flies*

CRISPR null flies were generated as described in FlyCripsr (https://flycrispr.org). Two guide RNAs (gRNA), one in exon 1 and another in exon 2 of the *bruno* gene, were designed (5'gRNA-CGGAGAAAUCGAAAAUCAUG and 3'gRNA-CGGCGAGAAGGAACCGGAUC) and cloned into an empty pU6-gRNA vector independently. Homology arms of 1kb each were cloned into the pHD-scarless dsRED donor vector. A mixture of gRNA plasmid and donor plasmid was injected into of *w*$^{1118}$*;*

*PBac{y[+mDint2]=vas-Cas9}VK00027* embryos. dsRED positive flies were selected and balanced with *CyO* and used as an *aret* knock out line.

*PhiC31 integrase-mediated site-specific insertion*

CDS of Bruno FL (1-604) and ΔN (147-604) were tagged at the C-terminus with mEGFP using the Gateway cloning system (Invitrogen) in an intermediate pActin5c vector. Subsequently the EGFP-tagged sequence was sub-cloned into the UASp-*attB* vector and injected into VK-33 (y*[1] M{vas-int.Dm}ZH-2A w[*]; PBac{y[+]-attP-3B}VK00033*) embryos for site-specific insertion into the *attP* landing site on chromosome III. Selection for positive transformants was based on eye color (*mini-white* gene). The transgene was balanced by *Tm3Ser* and the *PhiC31* integrase was removed in the final fly lines. For Hrp48 truncations, EGFP tagging was done at the N-terminus of FL and ΔC (1-205) using the Gateway cloning system (Invitrogen) in an intermediate pActin5c vector and then sub-cloned into the UASp-*attB* vector. Injection was done in embryos of VK-18 (*vas-phi-ZH2A, PBac{y[+]-attP-9A}VK00018*) flies for site-specific insertion into the *attP* landing site on chromosome II. The transgenes were balanced with *CyO*.

**Live imaging**

Ovaries of the desired genotype were dissected and mounted onto glass-bottom dishes in a 20 µl drop of Schneider's medium (with 10% fetal calf serum (FCS) and 200 µg/ml insulin) with an adjacent drop of Voltalef 10S oil. Individual egg chambers of desired stages were pulled under the oil with fine tungsten needles under a stereo microscope. Live imaging was done on an Zeiss LSM880 Airy Scan microscope (Airy Fast mode) with 40X/1.1 NA water immersion objective at room temperature.

***Ex vivo* treatments of whole ovaries**

*1,6-Hexanediol*: Ovaries of the desired genotype were dissected in PBS and incubated in Schneider's medium (with 10% FCS and 200 µg/ml insulin) with 5% 1,6-HD for 15 min at RT on a nutator. Water was used as a solvent control.

*Colchicine*: Ovaries of the desired genotype were dissected in PBS and incubated in Schneider's medium (with 10% FCS and 200 µg/ml insulin) with 100 µg/ml Colchicine for 2h at RT on a nutator. 100% ethanol was used as a solvent control. Live imaging was done after 90 min of colchicine treatment (100 µg/ml RT). Colchicine was present in the imaging medium.

**Protein expression in insect cells**

All plasmids are listed in Key Resources table. pFastBAC1 and pFastBAC-based pCoofy63 vector were used for cloning. pCoofy63 vector has a N-terminal 6xHis-SumoStar tag and a C-terminal Twin-Strep tag. Recombinant proteins Bruno, Hrp48 and PTB were expressed and purified from insect cells (Sf-21) using the baculovirus expression system to mimic the close-to-physiological state of the proteins. For generation of recombinant bacmid, pFastBAC1 or pFastBAC1-based pCoofy63 shuttle vector was transformed into DH10EmBacY *E. coli* competent cells via electroporation followed by blue-white screening to select for recombinant bacmid. This was followed by bacmid isolation and PCR-based verification. Sf-21 cells grown at a density of 0.5-1.0x$10^6$ cells/ml were transfected with recombinant bacmid using XtremeGene transfection reagent (Roche) and $V_0$ harvested after 72h of transfection. Virus amplification was carried out by infecting 25 ml Sf-21 cells with $V_0$ and $V_1$ harvested one day post proliferation arrest. Usually 1 L of uninfected Sf-21 cells (0.5-0.7x$10^6$ cell/ml) were infected with $V_1$ at a ratio of 1:100 and cells were harvested 72h after infection. Harvested cells were flash-frozen in liquid $N_2$ and stored at -80ºC.

**Protein purification**

*Bruno-EGFP, Bruno ∆N-EGFP, Hrp48-EGFP, & Hrp48 ∆C-EGFP*

Sf-21 cells expressing recombinant proteins were resuspended in lysis buffer (20 mM Tris-HCl pH 7.5, 500 mM NaCl, 1 mM EDTA supplemented with 0.01% TritonX-100, 1x tablet of Complete Mini Protease Inhibitor cocktail (Roche), 2 mM $MgCl_2$, Benzonase (Sigma)) for 10 min on ice for digestion of RNA/DNA by Benzonase nuclease followed by lysis using Microfluidizer. Lysate was centrifuged at 16000x g at 4ºC for 20 min to remove debris.

Proteins were C-terminally tagged with TwinStrep tag and affinity purification was done from the clarified lysate using 5ml StrepTrap HP column. The N-terminal 6xHis-Sumostar solubility tag was not cleaved during purification. Briefly, the clarified lysate was injected into the column, the bound proteins washed in 5-6 column volumes wash buffer (20 mM Tris-HCl pH 7.5, 500 mM NaCl, 1 mM EDTA) and finally eluted in Elution buffer (2.5 mM desthiobiotin (Sigma) in 20 mM Tris-HCl pH 7.5, 500 mM NaCl, 1 mM EDTA). Fractions were analysed in SDS-PAGE and desired fractions pooled in and dialysed overnight at 4ºC using a 12-14 kDa MWCO membrane (Spectrapor) in dialysis buffer (20 mM Tris-HCl pH 7.5, 500 mM NaCl) to remove EDTA and desthiobiotin. Post dialysis, the protein was concentrated to 5 ml and subjected to high resolution size exclusion chromatography in storage buffer (20 mM Tris-HCl pH 7.5, 300 mM NaCl, 2m M $MgCl_2$, 5% glycerol, 0.5 mM TCEP) using HiLoad 16/600 Superdex 200pg column. Fractions were analyzed in SDS-PAGE and desired fractions pooled and concentrated using concentrator with 50 kDa MWCO (Amicon). Droplet formation was checked during the purification to ensure that phase separation or aggregation does not occur during the concentration. Aliquots were flash-frozen and stored at -80ºC.

*mRFP-PTB*

mRFP-PTB was affinity purified by Ni-NTA chromatography. Briefly, Sf-21 cells were resuspended in Lysis buffer (20 mM Tris-HCl pH 7.5, 500 mM NaCl, 5% glycerol, 40 mM imidazole, supplemented with 0.01% TritonX-100, 1x tablet of Complete Mini

Protease Inhibitor cocktail (Roche), 2 mM $MgCl_2$, Benzonase (Sigma)) for 10 min on ice for digestion of RNA/DNA by Benzonase nuclease followed by lysis using Microfluidizer. Lysate was centrifuged at 16000x g at 4ºC for 20 min to remove debris. The clarified lysate was injected into a HisTrap HP column 5 ml, bound fractions were washed with 3-5 column volumes of wash buffer (20 mM Tris-HCl pH 7.5, 500 mM NaCl, 5% glycerol, 40 mM Imidazole) and eluted using a gradient of 40-600 mM Imidazole. Fractions were analysed in SDS-PAGE, desired fractions pooled and dialyzed overnight at 4ºC using a 12-14 kDa MWCO membrane (SpectraPor) in wash buffer to remove imidazole along with TEV protease (homemade in Protein Expression and Purification Core facility, EMBL) to cleave off the 6xHis tag. The untagged protein was separated from the 6xHis-tagged proteins and cleaved tags by a second round of Ni-NTA chromatography and the flow through (containing tag-cleaved protein) was collected, concentrated and subjected to high resolution size exclusion chromatography in storage buffer (20 mM Tris-HCl pH 7.5, 300 mM NaCl, 2 mM $MgCl_2$, 5% glycerol, 0.5 mM TCEP) using HiLoad 16/600 Superdex 200pg column. Fractions were analyzed by SDS-PAGE and desired fractions pooled and concentrated using a concentrator with 50 kDa MWCO (Amicon MERCK). Droplet formation was checked to ensure that phase separation or aggregation did not occur during the concentration steps. Aliquots were flash-frozen and stored at -80ºC.

All purifications were done using an Akta FPLC system (GE Life sciences). Protein extinction coefficients were calculated using ProtParam (Expasy) and concentrations were measured with diluted samples at 280 nm in a NanoDrop (Thermo Scientific).

**_In vitro_ transcription and fluorescent labelling of transcripts**

*In vitro* transcription (IVT) was performed using a MEGAscript T7 transcription kit (Invitrogen) according to the manufacturer’s instructions. Briefly, template for IVT was prepared by PCR using T7-forward primer and gene specific reverse primers. 200 ng template DNA was used for a 20 µl transcription reaction for 2-3h at 37ºC; template DNA was digested with Turbo DNAse and RNA was precipitated with LiCl and dissolved in ultrapure water (Invitrogen). For fluorescent labelling, the transcription reaction was spiked with 5-amino-allyl UTP (Biotium) at 1:4 (amino allylUTP: UTP). Fluorescent labelling was carried out with 3-fold molar excess of atto633 NHS-ester (Atto-Tec GmbH) in 0.1 M $NaHCO_3$ at RT for 2h, protected from light. RNA was precipitated at -20ºC with absolute ethanol and sodium acetate, pH 5.5, centrifuged at 16000x g 15 min 4ºC, followed by 2 washes with ice-cold 70% ethanol. RNA was dissolved in in ultrapure water (Invitrogen). Integrity of both the unlabeled and labelled transcripts was verified with SYBR Safe stain (473 nm), as well as by fluorescent gel imaging (635 nm) in a Typhoon biomolecular imager.

***In vitro* phase separation assays**

All *in vitro* phase separation experiments were carried out in assay buffer (20 mM Tris-HCl pH 7.5, 150 mM NaCl, 2 mM $MgCl_2$, 5% glycerol, 0.5 mM TCEP). No crowding agent was included, unless mentioned in figure descriptions. Protein concentrations were adjusted so that the final salt concentration was 150 mM NaCl in the assay.

Frozen aliquots of proteins were thawed and centrifuged at maximum speed to clear aggregates. For Bruno-EGFP and Hrp48-EGFP, the 6xHis-SumoStar tag was cleaved during the phase separation reaction using 1 U SumoStar protease (Life Sensors) per 20 µl reaction for 30 min at RT. *oskar* 3'UTR RNA (labelled or unlabeled) was added to the assay where specified. Details of individual experiments are indicated in respective figure schematics and legends. Reactions with all the components were assembled in Eppendorf tubes and immediately spotted on 96-well non-binding µclear plates (Greiner Bio-one), incubated up to 30 min for enzymatic tag removal and imaged using a Leica SP8 confocal microscope. For *in vitro* condensate ageing assays, the reactions were incubated in Eppendorf tubes for 30 min, followed by addition of RNA/proteins and then spotted onto 96-well plates for imaging.

**Electrophoretic Mobility Shift Assay (EMSA)**

EMSA was carried out as described previously (Besse et al., 2009). 50 nM of atto-633-labelled *oskar* 3'UTR (labeled as described above) was incubated with increasing concentrations of indicated proteins for 20 min at RT in assay buffer (20 mM Tris-HCl pH 7.5, 150 mM NaCl, 2 mM $MgCl_2$, 5% glycerol, 0.5 mM TCEP). The reactions were resolved on a 0.8% agarose gel in 0.5X TBE run at 100V constant current at 4ºC. Imaging of the gel was done by fluorescent gel imaging (635 nm) in a Typhoon biomolecular imager.

**Single molecule Fluorescent in situ hybridization (smFISH)**

The protocol used for smFISH has been described in detail in Gaspar et al., 2017.

*Labelling of $NH_2$-ddUTP:* Atto633-NHS-ester or Atto565-NHS-ester (Atto-Tec GmbH) was reconstituted with anhydrous DMSO in a desiccation chamber to 20 mM final concentration. Conjugation of Amino-11-ddUTP (NH2-ddUTP, Lumiprobe) with NHS-esters was done using a 2-fold molar excess of dye–NHS-ester in the presence of 0.1 M NaHCO3 pH 8.3 for 2h at RT protected from light. The reaction was quenched by adding 1 M Tris HCl pH 7.4 to 10 mM final concentration and the concentration adjusted to 5 mM with nuclease free water.

*Probe labelling* : Probe sequences are described in Table S1. Non-overlapping DNA oligos 18-22 nt (Sigma) long were selected using the smFISHprobe_finder.R script (Gaspar et al., 2018) and reconstituted to 250 µM with nuclease-free water. An equimolar mixture of a probe set was enzymatically conjugated to fluorescently labelled ddUTP (atto633 or atto565) using Terminal Deoxynucleotidyl Transferase (Thermo Scientific). Labelled oligos were precipitated with absolute ethanol, sodium acetate pH 5.5 and linear acrylamide and reconstituted with nuclease free water.

*Hybridization* : Freshly dissected ovaries were immediately fixed in 2% para-formaldehyde (PFA) in PBS with 0.05% TritonX-100 at RT for 20 min. The fixed ovaries were rinsed once and washed for 10 min with PBS containing 0.1% TritonX-100, followed by prehybridization at 42ºC for 10 min with shaking in hybridization buffer (300 mM NaCl, 30 mM sodium citrate pH 7.0, 15 v/v% ethylene carbonate, 1 mM EDTA, 50 µg/mL heparin, 100 µg/mL salmon testes DNA, 1% Triton X-100). 50 µL of prewarmed probe mixture (2.5 nM per individual oligonucleotide) was added and hybridization performed for 2-3h at 42°C. Excess probe was washed with 2 washes with hybridization buffer at 42ºC for 20 min each, followed by a final wash in PBS containing 0.1% TritonX-100. Ovaries were mounted in mounting medium (80% glycerol, 2% propyl gallate).

**Immunostaining and western blotting**

For immunostaining, freshly dissected ovaries were fixed in 4% PFA in PBS for 20 min, extracted in permeabilization buffer (1% TritonX-100 in PBS) and blocked in blocking buffer (0.5% BSA, 0.3% TritonX-100 in PBS) for 1h. Incubation with primary antibodies was done overnight at 4ºC followed by three washes in wash buffer (0.1% TritonX-100 in PBS) 15 min each. Incubation with Alexa Fluor-secondary antibodies diluted in 10% goat serum in PBS was carried out for 2h at RT, followed by 3 washes in wash buffer for 15 min each. Nuclei were stained with DAPI (1:2500 in wash buffer). Samples were mounted in mounting media (80% glycerol, 2% propyl gallate).

For western blotting, an equal number of freshly dissected ovaries of desired genotypes was directly resuspended and lysed in 1x Laemmli buffer (Invitrogen) supplemented with 5% β-mercaptoethanol. After boiling at 95ºC for 10 min, samples were centrifuged at maximum speed to remove debris and equal volumes loaded for SDS-PAGE analysis in 4-12% NuPAGE pre-cast gels (Invitrogen). Following wet transfer to PVDF membrane (Millipore) for 2h at 4ºC, the membrane was blocked in

5% skimmed milk in TBST (TBS-0.1% Tween-20), incubated with primary antibody overnight at 4ºC followed by 3 washes in TBST 10 min each. Incubation with secondary antibody was done at RT for 1h followed by 3 washes and detection by chemiluminescence (BioRad).

**Pole cell staining**

10-15 virgin females of the desired genotype were mated with double the number of $w^{1118}$ males and maintained on yeast paste for 2-3 days at 25ºC before egg collection. 0-4h old eggs were collected on apple juice agar plates, dechorionated with 50% bleach for 2 min, and fixed in preheated fixation buffer (0.4% NaCl, 0.3% Triton X-100 in PBS) at 92°C for 30s. This was immediately followed by devitellinization by rigorous shaking in a 1:1 mix of heptane and methanol. Fixed embryos were stored in 100% methanol at -20°C. For pole cell staining, fixed embryos of the desired genotype from multiple collections were pooled, washed 3-5 times with 0.1% Triton X-100 in PBS, blocked with blocking buffer (0.5% BSA, 0.3% TritonX-100 in PBS) and incubated with anti-Vasa (1:500) primary antibody overnight at 4°C. This was followed by 3 washes in wash buffer, incubation with secondary antibody at RT for 1h. The nuclei were stained with nuclear stain 4′,6-diamidino-2-phenylindole (DAPI).

**Embryonic cuticle preparations**

10-15 virgin females of the desired genotype were mated with double the number of $w^{1118}$ males while being fed with yeast paste for 2-3 days at 25ºC. Prior to egg collection, flies were placed in cages and allowed to lay eggs overnight on apple juice agar plates. Next morning the plates were collected and the eggs aged for 24h at 25ºC. After collection, the eggs were dechorionated with 50% bleach for 2 min, washed extensively with water and transferred to glass slides. Excess water was removed and the samples were mounted in Hoyer's medium and Lactic acid (Sigma), covered with a cover slip and baked overnight at 65ºC. Imaging was performed on a bright field microscope.

**Transfection of S2R+ cells**

*Drosophila* Schneider cells (S2R+) were cultured in Schneider's Drosophila medium (Life Technologies) containing 10% FCS (Life Technologies) and 1% Penicillin-Streptomycin. Transient transfection of cells was carried out using Effectene Transfection reagent (Qiagen) using manufacturer's protocol. Cells were transfected with 200ng plasmid in 24 well plates. After 24 hours cells were re-plated on Concanavalin A-coated cover slips for 1 hour prior to fixation with 4% PFA. Post-fixation cover slips were washed with 0.1% TritonX-100 containing PBS, mounted using Shandon ImmuMount (Thermo Scientific) and imaged with a water immersion objective.

**Image acquisition**

For high resolution image acquisitions of egg chambers, laser scanning confocal microscopy was carried out using a Leica TCS SP8 microscope with a HC PL APO 63x/1.30 Glycerol CORR CS2 glycerol-immersion objective. Images were acquired in the Leica Lightning mode with adaptive deconvolution. Low resolution imaging was done with a 20X/0.75 Air objective. For *in vitro* droplet experiments, images were acquired with an HC PL APO 40x/1.10 W CORR CS2 water-immersion objective without subsequent deconvolution.

3D STED microscopy was performed on a Leica STED 3X microscope equipped with a HC PL APO CS2 93X/1.30 Glycerol-immersion objective. Atto-633 was excited at 633nm with a white-light laser (WLL) and STED was performed at 775nm. Images were collected in line averaging mode (16 lines) and the pinhole was set to 1.0 Airy units. For XZ-scan, a pixel size of 40nmx12nm was used. 3D volumes were acquired with a voxel size of 40 nm x 40 nm x 60 nm. Acquired STED images were deconvolved with Huygens Professional (SVI) prior to analysis.

**Fluorescence Recovery After Photobleaching**

For Fluorescence Recovery After Photobleaching (FRAP) recordings, *in vitro* reconstituted condensates (assembled with Bruno-EGFP /Hrp48-EGFP/hFUS-EGFP) were excited with the 488 nm line of a white light laser on a Leica SP8 laser scanning confocal using a HC PL APO 40x/1.10 W CORR CS2 water-immersion objective. A circular region of interest of 1.4 -1.5 $\mu m^2$ was selected within the condensate and bleaching was carried out with 100% laser power of the same laser line. Three images were acquired prior to bleaching, following which fluorescence intensity was recorded for up to 100s at a frame interval of 0.35s. Movies were analyzed in Fiji and analysis was done using FRAPAnalyzer using double normalization and scaling the dynamic range from 0 to 1. Recovery curves were fitted with single exponential recovery using the formula: $FRAP(t)= I_0 + I_1 \cdot [1 - e^{-(t-tbleach)/\tau}]$

where, $I_0$ = normalized intensity just after bleach, $I_1$ = dynamic range of recovery, $\boldsymbol{\tau}$ = half-life of recovery. The immobile fraction was calculated as $(1- I_0- I_1)/(1- I_0)$.

**Quantification of absolute protein concentrations in granules *in vivo***

For calculating absolute concentrations per granule, Bruno and PTB EGFP-trap fly lines tagged with EGFP at the endogenous locus were used. A calibration curve was generated using bacterially expressed recombinant EGFP. Freshly dissected ovaries were fixed with 4% PFA for 15 min followed by washes with PBS containing 0.1% Triton X-100. Fixed egg chambers were separated in recombinant EGFP storage buffer (20 mM Tris-HCl pH 7.5, 150 mM NaCl, 2 mM $MgCl_2$, 5% glycerol, 0.5 mM TCEP) in 96-well non-binding µclear plates (Greiner Bio-one) and imaged with a Leica SP8 confocal microscope using a 40X HC PL APO 40x/1.10 W CORR CS2 water-immersion objective. Imaging was done in a cortical plane to resolve individual granules near the posterior cortex. A range of recombinant EGFP concentrations was imaged under identical optical settings at identical height from the cover slip surface. For calculations, single granules were marked with a ROI and concentrations estimated from a calibration curve of EGFP intensities.

**Quantification of *oskar* RNA concentrations in granules *in vivo***

*oskar* smFISH was carried out on wild type ($w^{1118}$) ovaries and egg chambers were imaged with a Leica SP8 using a HC PL APO 63x/1.30 Glycerol CORR CS2 glycerol-immersion objective in photon-counting mode and deconvolved using Leica Lightning adaptive deconvolution. Particles were segmented in xsPT Fiji plugin and intensities quantified. mRNA copy number was calculated by fitting multiple Gaussian functions to the corresponding signal intensity distributions taken from the nurse cells in R studio. The μ value of Gaussian fit that described the largest portion of the distribution in the nurse cells was taken as the signal intensity of a unit (the intensity of a single mRNA molecule). The volume of granules as obtained from 3D STED volume imaging was then used to calculate absolute mRNA concentration per granule.

**Cryo-electron tomography**

*Screening condensates on TEM grids*

All assay conditions were checked with light microscopy on standard non-binding glass surface. Prior to vitrification, samples were also screened on EM grids to ensure stability of condensates on the grid material. Quantifoil R2/1 Cu 200 mesh holey Carbon grids were glow-discharged for 45 s. 4 μl of sample was spotted on the grid, incubated for 1 min, followed by spiking of glutaraldehyde (fixative) to final concentration of 0.05% and imaged with a Zeiss Axiovert wide-field microscope after 30 s.

*Vitrification by plunge freezing*

After assessing condensate stability on grids, identical conditions were used for sample spotting on the grids and proceeded to plunge-freezing. Grids were blotted from both sides for 2s with 0 blot force, followed by a drain time of 2s and immediately plunged into liquid ethane at liquid nitrogen temperature using a Vitrobot Mark 4 (FEI Company/ThermoFisher Scientific, Eindhoven, Netherlands) with the chamber set at 22ºC, 90% humidity. The frozen grids were in liquid nitrogen until further processing.

*Cryo-Electron tomography*

Cryo-electron tomography was carried out on a 300 kV (FEI Company/ThermoFisher Scientific, Eindhoven, Netherlands) Titan Krios microscope equipped with a field-emission gun, a Quantum post-column energy filter (Gatan, Pleasanton, CA, USA), a K2 direct detector camera (Gatan) and a Volta phase plate (FEI Company/ThermoFisher Scientific, Eindhoven, Netherlands). Data recording was done in dose fractionation mode using SerialEM software v3.7.2 (Mastronarde, 2005). Tilt-series were collected using a dose symmetric scheme (Hagen et al., 2017) and a Volta phase plate (Danev et al., 2014) in nano-probe mode, pixel size at the specimen level of 2.12 Å, 3-4 µm defocus, tilt increment of 2º at a constant dose of 2.1 $e^{-}$/ $Å^{2}$ for all tilts. Motion correction and Contrast transfer function (CTF) correction was performed using WARP (Tegunov and Cramer, 2019). Tilt-series alignment and tomographic reconstructions were done using the IMOD software package, version 4.9.4 (Kremer et al., 1996) with patch-tracking. Aligned images were binned to the final pixel size of 8.51 Å. For tomographic reconstruction by back-projection, the radial filter options were set at default values (cut off, 0.35; fall off, 0.05). The reconstructed tomograms were filtered with a Gaussian filter of radius of 3 pixels using the TOM package implemented in Matlab (Mathworks).

**Image analysis**

*Analysis of STED data*

For quantification of 3D STED volumes, the volumes were analysed and segmented using Imaris (BitPlane) and statistics of the segmented particles used for plotting of volumes, intensities and aspect ratios.

*Fiji based analysis*

For all image representations in the figures, Fiji was used. Intensity-based thresholding, histogram generation & intensity line profile was done with Fiji. Maximum intensity Z-projections were also done with Fiji. For calculation of granule size, intensity-based thresholding was done in Fiji followed by particle analysis.

*Cortical Analysis*

For cortical analysis CortAnalysis Fiji plugin was used (Gaspar et al., 2014).

*Quantification of Oskar protein levels*

For quantification of Oskar protein signal in Figure 7 and related Supplementary figures, the region containing the signal was manually selected using a polygon tool and the integrated density quantified.

**Statistical analysis**

Statistical analyses were performed using GraphPad Prism8. P-values were calculated using unpaired two-tailed t-test. In the figures, * = p< 0.05, ** = p< 0.01, *** = p< 0.001, ****= p< 0.0001.
